# Predicting distributed working memory activity in a large-scale mouse brain: the importance of the cell type-specific connectome

**DOI:** 10.1101/2022.12.05.519094

**Authors:** Xingyu Ding, Sean Froudist-Walsh, Jorge Jaramillo, Junjie Jiang, Xiao-Jing Wang

## Abstract

Recent advances in connectome and neurophysiology make it possible to probe whole-brain mechanisms of cognition and behavior. We developed a large-scale model of the mouse multiregional brain for a cardinal cognitive function called working memory, the brain’s ability to internally hold and process information without sensory input. The model is built on mesoscopic connectome data for inter-areal cortical connections and endowed with a macroscopic gradient of measured parvalbumin-expressing interneuron density. We found that working memory coding is distributed yet exhibits modularity; the spatial pattern of mnemonic representation is determined by long-range cell type-specific targeting and density of cell classes. Cell type-specific graph measures predict the activity patterns and a core subnetwork for memory maintenance. The model shows numerous self-sustained internal states (each engaging a distinct subset of areas). This work provides a framework to interpret large-scale recordings of brain activity during cognition, while highlighting the need for cell type-specific connectomics.

## Introduction

In contrast to our substantial knowledge of local neural computation, such as orientation selectivity in the primary visual cortex or the spatial map of grid cells in the medial entorhinal cortex, much less is understood about distributed processes in multiple interacting brain regions underlying cognition and behavior. This has recently begun to change, as advances in new technologies enable neuroscientists to probe neural activity at single-cell resolution and on a large-scale by electrical recording or calcium imaging of behaving animals (Jun et al. 2017; Steinmetz et al. 2019; Stringer et al. 2019; Musall et al. 2019; Steinmetz et al. 2021), ushering in a new era of neuroscience investigating distributed neural dynamics and brain functions (Wang 2022).

To be specific, consider a core cognitive function called working memory, the ability to temporally maintain information in mind without external stimulation (Baddeley 2012). Working memory has long been studied in neurophysiology using delay-dependent tasks, where stimulus-specific information must be stored in working memory across a short time period between a sensory input and a memory-guided behavioral response (Fuster and Alexander 1971; Funahashi et al. 1989; Goldman-Rakic 1995; Wang 2001). Delay-period mnemonic persistent neural activity has been observed in multiple brain regions, suggesting distributed working memory representation (Suzuki and Gottlieb 2013; Leavitt et al. 2017; Christophel et al. 2017; Xu 2017; Dotson et al. 2018). Connectome-based computational models of the macaque cortex found that working memory activity depends on interareal connectivity (Murray et al. 2017; Jaramillo et al. 2019), macroscopic gradients of synaptic excitation (Wang 2020; Mejias and Wang 2022) and dopamine modulation (Froudist-Walsh et al. 2021).

Mnemonic neural activity during a delay period is also distributed in the mouse brain (Liu et al. 2014; Schmitt et al. 2017; Guo et al. 2017; Bolkan et al. 2017; Gilad et al. 2018). The new recording and imaging techniques as well as optogenetic methods for causal analysis (Yizhar et al. 2011), that are widely applicable to behaving mice, hold promise for elucidating the circuit mechanism of distributed brain functions in rodents. Recurrent synaptic excitation represents a neural basis for the maintenance of persistent neural firing (Goldman-Rakic 1995; D. J. Amit 1995; Wang 2021). In the monkey cortex, the number of spines (sites of excitatory synapses) per pyramidal cell increases along the cortical hierarchy, consistent with the idea that mnemonic persistent activity in association cortical areas including the prefrontal cortex is sustained by recurrent excitation stronger than in early sensory areas. Such a macroscopic gradient is lacking in the mouse cortex (Gilman et al. 2017; Ballesteros-Yáñez et al. 2010), raising the possibility that the brain mechanism for distributed working memory representations may be fundamentally different between mice and monkeys.

In this paper we report a cortical mechanism of distributed working memory that does not depend on a gradient of synaptic excitation. We developed an anatomically-based model of the mouse brain for working memory, built on the recently available mesoscopic connectivity data of the mouse thalamocortical system (Oh et al. 2014; Gămănuţ et al. 2018; Harris et al. 2019; Kim et al. 2017). Our model is validated by capturing large-scale neural activity observed in recent mouse experiments (Guo et al. 2017; Gilad et al. 2018). Using this model, we found that a decreasing gradient of synaptic inhibition mediated by parvalbumin (PV) positive GABAergic cells (Kim et al. 2017; Fulcher et al. 2019; Wang 2020) and long-range excitatory connections shape the distributed pattern of working memory representation. Moreover, the engagement of inhibition through local and long range projections determines the stability of the local circuits, further emphasizing the importance of inhibitory circuits. A focus of this work is to examine whether anatomical connectivity can predict the emergent large-scale neural activity pattern underlying working memory. Interestingly, traditional graph-theory measures of inter-areal connections, which ignore cell types of projection targets, are uncorrelated with activity patterns. We propose new cell type-specific graph theory measures to overcome this problem, and differentiate contributions of cortical areas in terms of their distinct role in loading, maintaining, and reading out the content of working memory. Through computer-simulated perturbations akin to optogenetic inactivations, a core subnetwork was uncovered for the generation of persistent activity. This core subnetwork can be predicted based on the cell type-specific interareal connectivity, highlighting the necessity of knowing the cell type targets of interareal connections in order to relate anatomy with physiology and behavior. This work provides a computational and theoretical platform for cross-scale understanding of cognitive processes across the mouse cortex.

## Results

### A decreasing gradient of PV interneuron density from sensory to association cortex

Our large-scale circuit model of the mouse cortex uses inter-areal connectivity provided by anatomical data within the 43-area parcellation in the common coordinate framework v3 atlas (Oh et al. 2014) (Fig. 1A, Fig. 1 - supplement 1A). The model is endowed with area-to-area variation of parvalbumin-expressing interneurons (PV) in the form of a gradient measured from the qBrain mapping platform (Fig. 1 - supplement 1B) (Kim et al. 2017). The PV cell density (the number of PV cells per unit volume) is divided by the total neuron density, to give the PV cell fraction, which better reflects the expected amount of synaptic inhibition mediated by PV neurons (Fig. 1B-C, neuron density is shown in Fig. 1 - supplement 1C). Cortical areas display a hierarchy defined by mesoscopic connectome data acquired using anterograde fluorescent tracers (Oh et al. 2014) (Fig. 1D-E). In Fig. 1F, the PV cell fraction is plotted as a function of the cortical hierarchy, which shows a moderate negative correlation between the two. Therefore, primary sensory areas have a higher density of PV interneurons than association areas, although the gradient of PV interneurons does not align perfectly with the cortical hierarchy.

**Figure 1.**
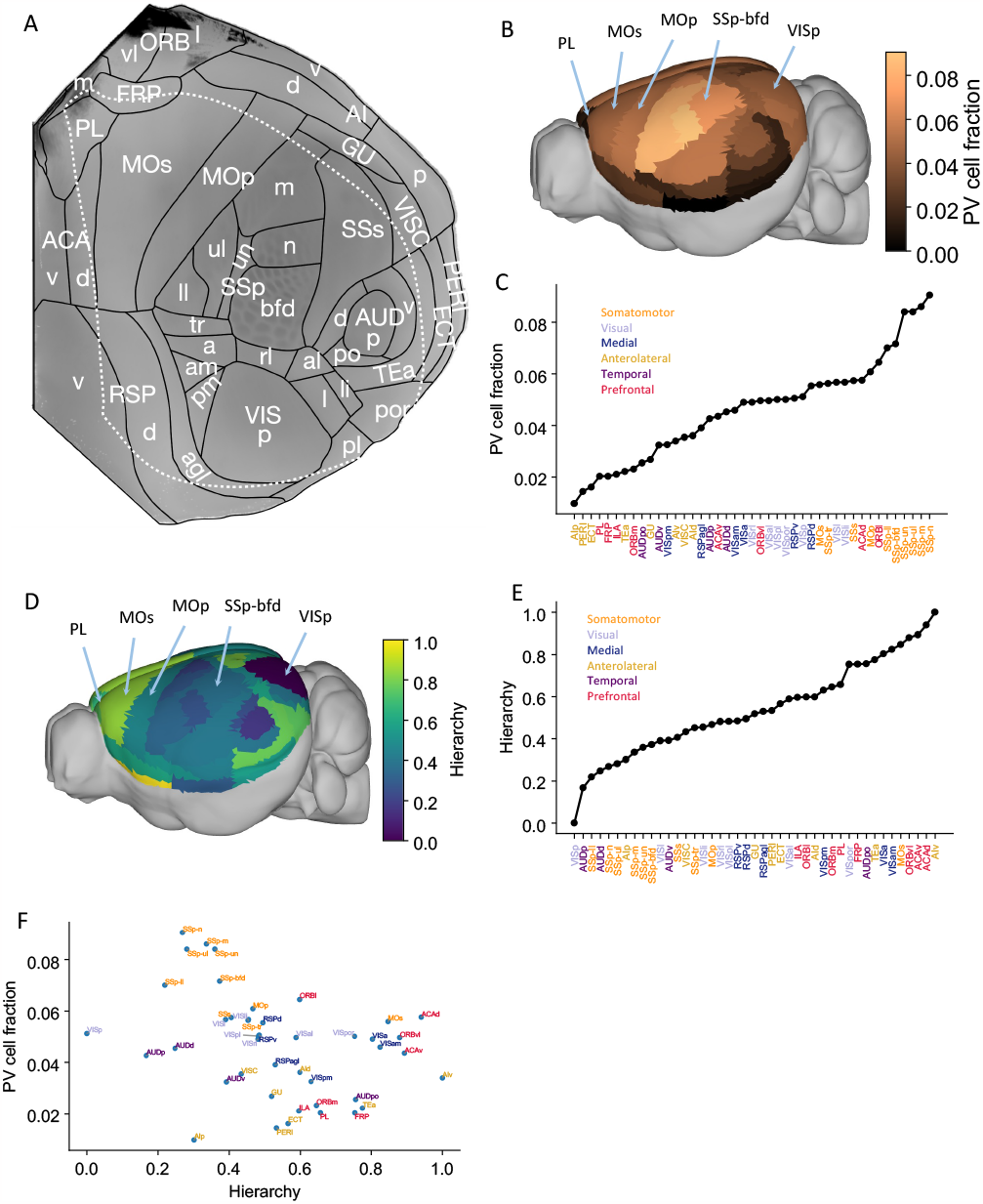
Anatomical basis of the multi-regional mouse cortical model. (A). Flattened view of mouse cortical areas. Figure adapted from (Harris et al. 2019). (B). Normalized PV cell fraction for each brain area, visualized on a 3d surface of the mouse brain. Five areas are highlighted : VISp, Primary somatosensory area, barrel field (SSp-bfd), primary motor (MOp), MOs and PL. (C). The PV cell fraction for each cortical area, ordered. Each area belongs to one of five modules, shown in color. (Harris et al. 2019). (D). Hierarchical position for each area on a 3d brain surface. Five areas are highlighted as in (B), and color represents the hierarchy position. (E). Hierarchical positions for each cortical area. The hierarchical position is normalized and the hierarchical position of VISp is set to be 0. As in C), the colors represent the module that an area belongs to. (F). Correlation between PV cell fraction and hierarchy (Pearson correlation coefficient r = −0.35, p < 0.05).

### A whole-mouse cortex model with a gradient of interneurons

In our model, each cortical area is described by a local circuit (Fig. 2A), using a mean-field reduction (Wong and Wang 2006) of a spiking neural network (Wang 2002). We use a version of this model that has two excitatory neural pools selective for different stimuli and a shared inhibitory neural pool to describe each cortical area. The model makes the following assumptions. First, local inhibitory strength is proportional to PV interneuron density across the cortex. Second, the inter-areal long-range connection matrix is given by the anterograde tracing data (Oh et al. 2014; Knox et al. 2018; Wang et al. 2020). Third, targeting is biased onto inhibitory cells for top-down compared with bottom-up projections. Therefore, feedforward connections have a greater net excitatory effect than feedback connections, which is referred to as counterstream inhibitory bias (CIB) (Mejias and Wang 2022; Javadzadeh and Hofer 2022; Wang 2022). Briefly, we assume that long-range connections are scaled by a coefficient that is based on the hierarchy of the source and target areas. According to the CIB assumption, long-range connections to inhibitory neurons are stronger for feedback connections and weaker for feedforward connections, while the opposite holds for long range connections to excitatory neurons.

**Figure 2.**
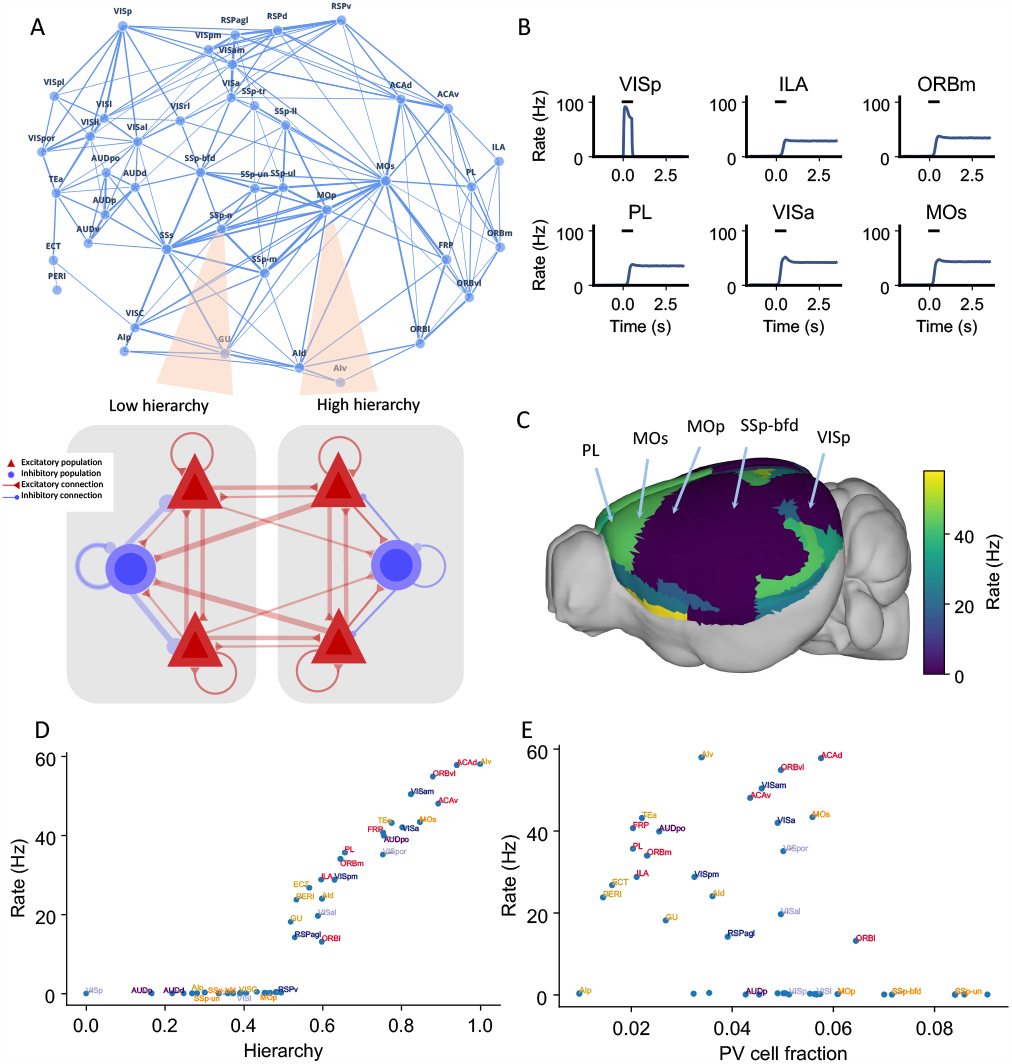
Distributed working memory activity depends on the gradient of PV interneurons and the cortical hierarchy. (A). Model design of the large-scale model for distributed working memory. Top, connectivity map of the cortical network. Each node corresponds to a cortical area and an edge is a connection, where the thickness of the edge represents the strength of the connection. Only strong connections are shown (without directionality for the sake of clarity). Bottom, local and long-range circuit design. Each local circuit contains two excitatory populations (red), each selective to a particular stimulus and one inhibitory population (blue). Long-range connections are scaled by mesoscopic connectivity strength (Oh et al. 2014) and follows counterstream inhibitory bias (CIB) (Mejias and Wang 2022). (B). The activity of 6 selected areas during a working memory task is shown. A visual input of 500ms is applied to area VISp, which propagates to the rest of the large-scale network. (C). Delay period firing rate for each area on a 3d brain surface. Similar to Fig. 1B, the positions of 5 areas are labeled. (D). Delay-period firing rate is positively correlated with cortical hierarchy (r = 0.91, p < 0.05). (E). Delay-period firing rate is negatively correlated with PV cell fraction (r = −0.43, p < 0.05).

### Distributed working memory activity depends on the gradient of inhibitory neurons and the cortical hierarchy

We simulated the large-scale network to perform a simple visual delayed response task that requires one of two stimuli to be held in working memory. We shall first consider the case in which the strength of local recurrent excitation is insufficient to generate persistent activity when parcellated areas are disconnected from each other. Consequently, the observed distributed mnemonic representation must depend on long-range interareal excitatory connection loops. Later in the paper we will discuss the network model behavior when some local areas are capable of sustained persistent firing in isolation.

The main question is: when distributed persistent activity emerges after a transient visual input is presented to the primary visual cortex (VISp), what determines the spatial pattern of working memory representation? After we remove the external stimulus, the firing rate in area VISp decreases rapidly to baseline. Neural activity propagates throughout the cortex after stimulus offset (Fig. 2B). Neural activities in the higher visual cortical areas (e.g. VISrl and VISpl) show similar dynamics to VISp. In stark contrast, many frontal and lateral areas (including prelimbic (PL), infralimbic (ILA), secondary motor (MOs) and ventral agranular insula (AIv) areas) sustained a high firing rate during the delay period (Fig. 2B). Areas that are higher in the cortical hierarchy show elevated activity during the delay period (Fig. 2C). This persistent firing rate could last for more than 10 seconds and is a stable attractor state of the network (Inagaki et al. 2019).

The cortical hierarchy and PV fraction predict the delay period firing rate of each cortical area (Fig. 2C-E). Thus the activity pattern of distributed working memory depends on both local and large-scale anatomy. The delay activity pattern has a stronger correlation with hierarchy (r = 0.91) than with the PV fraction (r = −0.43). The long-range connections thus play a predominant important role in defining the persistent activity pattern.

Activity in early sensory areas such as VISp displays a rigorous response to the transient input but returns to a low firing state after stimulus withdrawal. In contrast, many frontal areas show strong persistent activity. When the delay period firing rates are plotted versus hierarchy, we observe a gap in the distribution of persistent activity (Fig. 2D) that marks an abrupt transition in the cortical space. This leads to the emergence of a subnetwork of areas capable of working memory representations.

We also used our circuit model to simulate delayed response tasks with different sensory modalities (Fig. 2 - supplement 1), by stimulating primary somatosensory area SSp-bfd and primary auditory area AUDp. The pattern of delay period firing rates for these sensory modalities is similar to the results obtained for the visual task: sensory areas show transient activity, while frontal and lateral areas show persistent activity after stimulus withdrawal. Moreover, the cortical hierarchy could predict the delay period firing rate of each cortical area well (r = 0.89, p < 0.05), while the PV cell fraction could also predict the delay period firing rate of each cortical area with a smaller correlation coefficient (r = *−*0.4, p < 0.05). Our model thus predicts that working memory may share common activation patterns across sensory modalities, which is partially supported by cortical recordings during a memory-guided response task (Inagaki et al. 2018).

We explored the potential contributions of PV gradients and CIB in determining spatially-patterned activity across the cortex. To evaluate the importance of the PV gradient, we replaced the PV gradient across areas with a constant value (Fig. 3A(ii)). As compared to the model with a PV gradient (Fig. 3A(i)), we found that, during the delay period, the number of cortical areas displaying persistent activity is diminished, but the abrupt transition in delay period firing rates remains. This quantitative difference depends on the constant value used to scale inhibition from PV cells across areas (Fig. 3 - supplement 1A, 1B). Next, we performed the analogous manipulation on the CIB by scaling feedforward and feedback projections with a constant value across areas, thus effectively removing the CIB. In this case, the firing rate of both sensory and association areas exhibit high firing rates during the delay period (Fig. 3A(iii)). Thus, CIB may be particularly important in determining which areas exhibit persistent activity.

**Figure 3.**
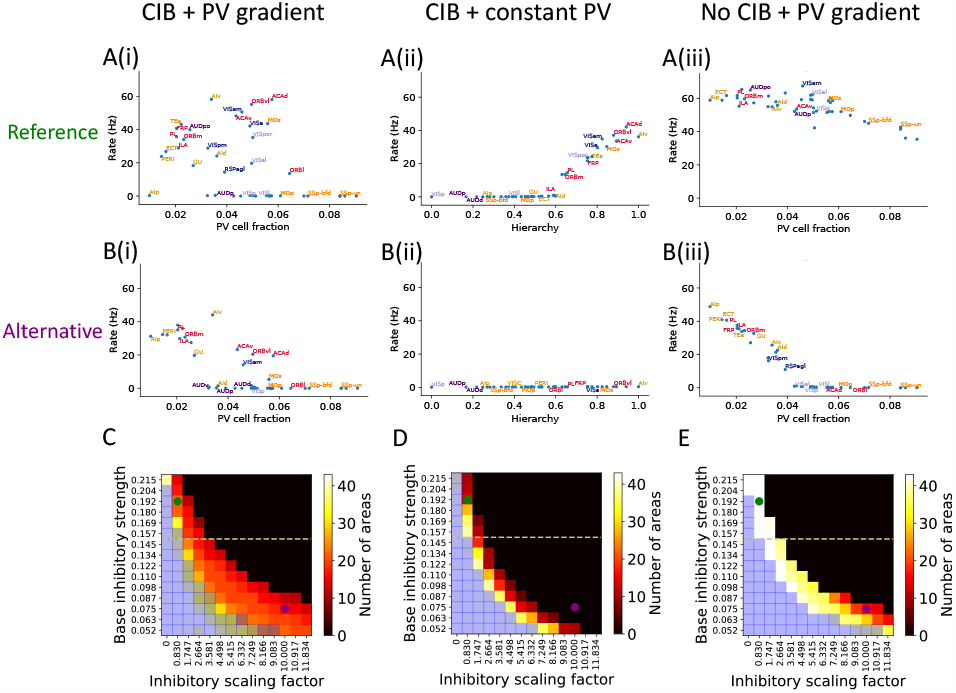
The role of PV inhibitory gradient and hierarchy-based counter inhibitory bias (CIB) in determining persistent activity patterns in the cortical network. (A(i)). Delay firing rate as a function of PV cell fraction with both CIB and PV gradient present (r =−0.42, p<0.05). This figure panel is the same as Fig. 2E. (A(ii)). Delay firing rate as a function of hierarchy after removal of PV gradient (r= 0.85, p<0.05). (A(iii)). Delay firing rate as a function of PV cell fraction after removal of CIB (r =−0.74, p<0.05). (B(i)). Delay firing rate as a function of PV cell fraction with both CIB and PV gradient present, in the alternative regime (r =−0.7, p<0.05). (B(ii)). Delay firing rate as a function of hierarchy after removal of PV gradient, in the alternative regime (r = 0.95, p<0.05). (B(iii)). Delay firing rate as a function of PV cell fraction after removal of CIB, in the alternative regime (r =−0.84, p<0.05). (C)-(E). Number of areas showing persistent activity (color coded) as a function of the local inhibitory gradient (g_EI,scaling_, X axis) and the base value of the local inhibitory gradient (g_EI,0_, Y axis) for the following scenarios: (C) CIB and PV gradient, (D) with PV gradient replaced by a constant value, and (E) with CIB replaced by a constant value. The reference regime is located at the top left corner of the heatmap (green dot) and corresponds to A(i)-A(iii), while the alternative regime is located at the lower right corner (purple dot) and corresponds to B(i)-B(iii). The yellow dashed lines separate parameters sets for which none of the areas show ‘independent’ persistent activity (above the line) from parameter sets for which some areas are capable of maintaining persistent activity without input from other areas (below the line). Blue shaded squares in the heatmap mark the absence of a stable baseline.

To further explore the model parameter space and better understand the interplay between PV gradient and CIB, we systematically varied two critical model parameters: i) the base local inhibitory weight *g*_*EI*,0_ onto excitatory neurons, which sets the minimal inhibition for each cortical area and ii) the scaling factor *g*_*EI,scaling*_, which refers to how strongly the PV gradient is reflected in the inhibitory weights. We created heatmaps that show the number of areas with persistent activity during the delay period as a function of these parameters: in Fig. 3C, we simulate the network with both CIB and PV gradient, while in Fig. 3D and Fig. 3E we simulate networks when PV gradient or CIB is removed, respectively. In each of these networks, we identify two regimes based on specific values for *g*_*EI*,0_ and *g*_*EI,scaling*_: a reference regime (used throughout the rest of the paper) and an alternative regime.

If we remove the PV gradient in the alternative parameter regime, persistent activity is lost (Fig. 3B(ii)). In contrast, if we remove CIB the model still exhibits an abrupt transition in firing rate activity (Fig. 3B(iii)). In this regime, a strong correlation and piece-wise linear relationship between firing rate and PV cell fraction was uncovered that did not exist when CIB was present. This observation led to a model prediction: if PV cell fraction is not strongly correlated with delay firing rate across cortical areas (e.g., Fig. 3A(i) or Fig. 3B(i)), this suggests the existence of a CIB mechanism at play. Importantly, the model without CIB exhibits the abrupt transition in delay-period firing rates provided it is in a regime where some areas exhibit ’independent’ persistent activity: persistent activity that is generated due to local recurrence and thus independent of long-range recurrent loops. The parameter regime where some areas exhibit ‘independent’ persistent activity is quantified by varying the base value of local inhibitory connections (Fig. 3 - supplement 1C). To conclude, the model results suggest that CIB may be present in a large-scale brain network if the PV cell fraction is not strongly correlated with the delay firing rate. Furthermore, CIB may be particularly important in the regime where local connections are not sufficient to sustain independent persistent activity.

Next, we evaluated the stability of the baseline state for the three conditions described above: i) original with PV gradient and CIB, ii) after removal of CIB, and iii) after removal of PV gradient. The heatmaps obtained after varying the base inhibitory strength and inhibitory scaling factor were qualitatively the same across the three conditions, as shown by the blue shaded squares in Fig. 3C-E. There are some regimes, such as the one depicted on the lower left corner, where all areas exhibit persistent activity and there is no stable baseline: a regime that is not biologically-realistic for a healthy brain. Thus, while PV and CIB shape the distribution of delay firing rates across cortical areas, they don’t qualitatively influence the system’s baseline stability. However, the inclusion of both PV gradient and CIB in the model (Fig. 3C) results in a more robust system, i.e., a far wider set of parameters can produce realistic persistent activity (Fig. 3D, 3E).

### Local and long-range projections modulate the stability of the baseline state in the cortex

The stability of the baseline state for any given cortical area may have contributions from local inhibition or from long-range projections that target local inhibitory circuits. We found that individual local networks without long-range connections are stable without local inhibition (Fig. 4A, see methods for theoretical calculation of stability in a local circuit). However, in the full network with long-range connections, setting either the long-range connections to inhibitory neurons or local inhibition to zero made the network’s baseline state unstable, and individual areas rose to a high firing rate (Fig. 4B). Thus, inhibition from local and long-range circuits contribute to the baseline stability of cortical areas.

**Figure 4.**
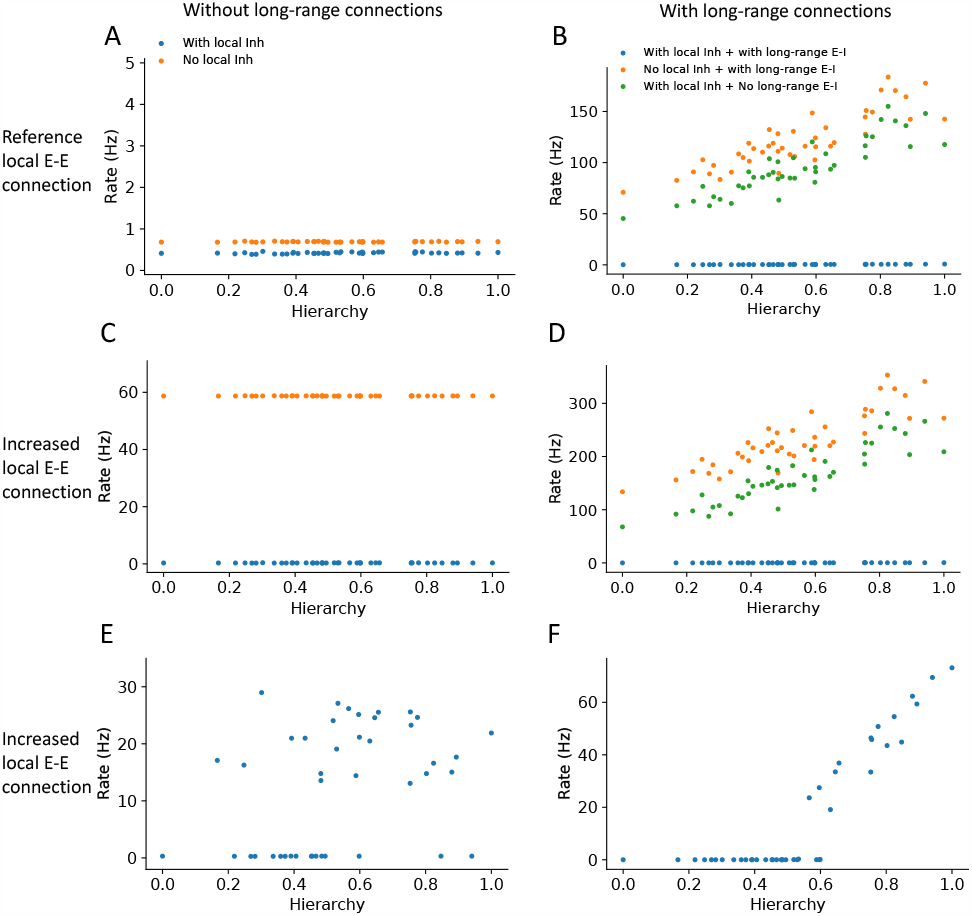
Local and long-range projections modulate the baseline stability of individual cortical areas. Steady state firing rates are shown as a function of hierarchy for different scenarios: (A) without long-range connections in the reference regime (g_E,self_ = 0.4nA, g_EI,0_ = 0.192nA), (B) with long-range connections in the reference regime (μ_EE_ = 0.1nA), (C) without long-range connections and increased local excitatory connections (g_E,self_ = 0.6nA, g_EI,0_ = 0.5nA), and (D) with long-range connections (increased long-range connections to excitatory neurons, μ_EE_ = 0.19nA) and increased local excitatory connections. (E) Firing rate as a function of hierarchy when external input given to each area, showing bistability for a subset of areas (parameters as in (C) and ‘with local inh’). (F) Firing rate as a function of hierarchy when external input is applied to area VISp (parameters as in (D)and ‘with local inh + with long-range E-I’).

Motivated by the results on stability, we investigated whether the large-scale network model operates in the inhibitory stabilized network (ISN) regime (Tsodyks et al. 1997; Sanzeni et al. 2020), whereby recurrent excitation is balanced by inhibition to maintain stability of the baseline state. First, we examined whether individual brain areas (i.e., without long range projections) may operate in this regime. We found a parameter set in which the baseline firing rate is stable only when local inhibition is intact: when inhibition is removed, the stable baseline state disappears, which suggests that the local circuits are ISNs (Fig. 4C and see stability analysis in the Methods section). In the full neural network with long-range connections, similar analysis as in Fig. 4B shows that the network becomes unstable if long-range projections onto inhibitory interneurons are removed. (Fig. 4D). Thus we propose that the network is also in a ’global’ inhibitory stabilized network (ISN) regime, whereby long-range connections to inhibitory neurons are necessary to maintain a stable baseline state. Second, Second, we examined whether the ISN regime is consistent with distributed working memory patterns in the cortex (Fig. 2). In the regime with increased local excitatory connections but without long-range projections, some local circuits could reach a high stable state when an external input is applied, demonstrating the bistability of those areas (Fig. 4E). When we considered the full network with long-range projections, the network exhibits a graded firing rate pattern after transient stimulation of VISp, showing that the interconnected ISN networks are compatible with bistability of a subset of cortical areas (Fig. 4F)

In summary, we have shown that distinct local and long-range inhibitory mechanisms shape the pattern of working memory activity and stability of the baseline state.

### Thalamocortical interactions maintain distributed persistent activity

To investigate how thalamocortical interactions affect the large-scale network dynamics, we designed a thalamocortical network similar to the cortical network (Fig. 5A). Several studies have shown that thalamic areas are also involved in the maintenance of working memory (Bolkan et al. 2017; Guo et al. 2017; Schmitt et al. 2017). However, the large-scale thalamocortical mechanisms underlying memory maintenance are unknown. We set the strength of connections between the thalamus and cortex using data from the Allen Institute (Oh et al. 2014) (Fig. 5 - supplement 1). All thalamocortical connections in the model are mediated by AMPA synapses. There are no recurrent connections in the thalamus within or across thalamic nuclei (Jones 2007). The effect of thalamic reticular nucleus neurons was included indirectly as a constant inhibitory current to all thalamic areas (Crabtree 2018; Hádinger et al. 2023). Similarly to cortical areas, the thalamus is organized along a measured hierarchy (Harris et al. 2019). For example, the dorsal part of the lateral geniculate nucleus (LGd) is lower than the cortical area VISp in the hierarchy, consistent with the fact that LGd sends feedforward inputs to VISp. Thalamocortical projections in the model are slightly more biased toward excitatory neurons in the target area if they are feedforward projections and towards inhibitory neurons if they are feedback.

**Figure 5.**
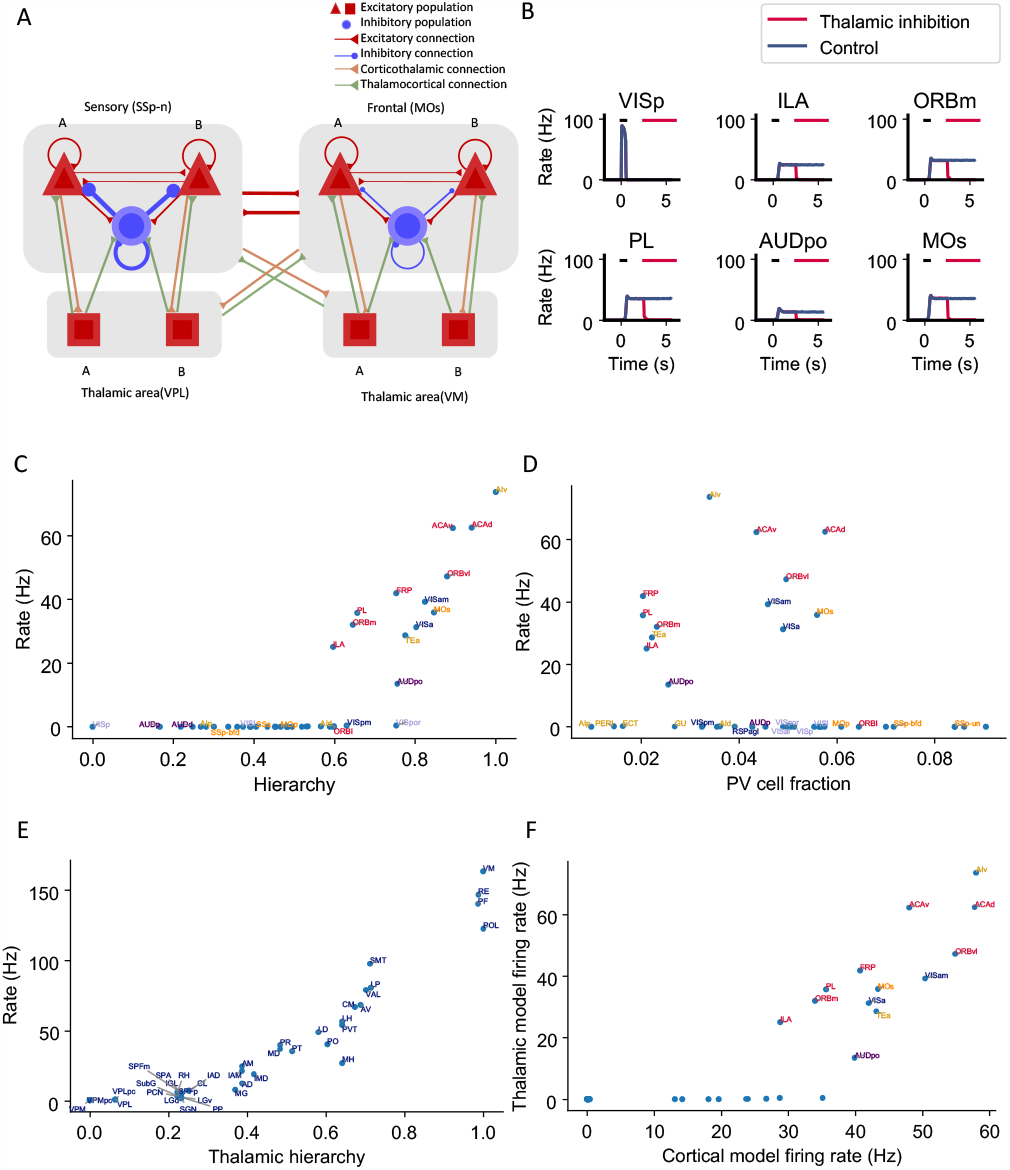
Thalamocortical interactions help maintain distributed persistent activity. (A). Model schematic of the thalamocortical network. The structure of the cortical component is the same as our default model in Fig. 2A, but with modified parameters. Each thalamic area includes two excitatory populations (red square) selective to different stimuli. Long range projections between thalamus and cortex also follow the counterstream inhibitory bias rule as in the cortex. Feedforward projections target excitatory neurons with stronger connections and inhibitory neurons with weaker connections; the opposite holds for feedback projections. (B). The activity of 6 sample cortical areas in a working memory task is shown during control (blue) and when thalamic areas are inhibited in the delay period (red). Black dashes represent the external stimulus applied to VISp. Red dashes represent external inhibitory input given to all thalamic areas. (C). Delay period firing rate of cortical areas in the thalamocortical network. The activity pattern has a positive correlation with cortical hierarchy (r = 0.78, p < 0.05). (D). Same as (C) but plotted against PV cell fraction. The activity pattern has a negative correlation with PV cell fraction, but it is not significant (r =−0.26, p = 0.09). (E). Delay firing rate of thalamic areas in thalamocortical network. The firing rate has a positive correlation with thalamic hierarchy (r = 0.94, p < 0.05). (F). Delay period firing rate of cortical areas in thalamocortical network has a positive correlation with delay firing rate of the same areas in a cortex-only model (r = 0.77, p < 0.05). Note that only the areas showing persistent activity in both models are considered for correlation analyses.

Here, we weakened the strength of cortical interareal connections as compared to the cortex model of Fig. 2. Now, persistent activity can still be generated (Fig. 5B, blue) but is maintained with the help of the thalamocortical loop, as observed experimentally (Guo et al. 2017). Indeed, in simulations where the thalamus was inactivated, the cortical network no longer showed sustained activity (Fig. 5B, red).

In the thalamocortical model, the delay activity pattern of the cortical areas correlates with the hierarchy, again with a gap in the firing rate separating the areas engaged in persistent activity from those that do not (Fig. 5B, Fig. 5C). Sensory areas show a low delay firing rate, and frontal areas show strong persistent firing. Unlike the cortex, the firing rate of thalamic areas continuously increases along the hierarchy (Fig. 5E). On the other hand, cortical dynamics in the thalamocortical and cortical models show many similarities. Early sensory areas do not show persistent activity in either model. Many frontal and lateral areas show persistent activity and there is an abrupt transition in cortical space in the thalamocortical model, like in the cortex only model. Quantitatively, the delay firing pattern of the cortical areas is correlated with the hierarchy and the PV fraction (Fig. 5C, Fig. 5D). Furthermore, the delay period firing rate of cortical areas in the thalamocortical model correlates well with the firing rate of the same areas in the cortical model (Fig. 5F). This comparison suggests that the cortical model captures most of the dynamical properties in the thalamocortical model; therefore in the following analyses, we will mainly focus on the cortex-only model for simplicity.

### Cell type-specific connectivity measures predict distributed persistent firing patterns

Structural connectivity constrains large-scale dynamics (Mejias and Wang 2022; Froudist-Walsh et al. 2021; Cabral et al. 2011). However, we found that standard graph theory measures could not predict the pattern of delay period firing across areas. There is no significant correlation between input strength and delay period firing rate (r = 0.25, p = 0.25, Fig. 6A(i), A(ii)) and input strength cannot predict which areas show persistent activity (prediction accuracy = 0.51, Fig. 6A(iii)). We hypothesized that this is because currently available connectomic data used in this model do not specify the type of neurons targeted by the long-range connections. For instance, when two areas are strongly connected with each other, such a loop would contribute to the maintenance of persistent activity if projections are mutually excitatory, but not if one of the two projections predominantly targets inhibitory PV cells. Therefore, cell type-specificity of interareal connections must be taken into account in order to relate the connectome with the whole-brain dynamics and function. To examine this possibility, we introduced a *cell type projection coefficient* (see *Calculation of network structure measures* in the Methods), which is smaller with a higher PV cell fraction in the target area (Fig. 6 - supplement 1). The cell type projection coefficient also takes cell type targets of long range connections into account, which, in our model, is quantified by counterstream inhibitory bias (CIB). As a result, the modified cell type-specific connectivity measures increase if the target area has a low density of PV interneurons and/or if long-range connections predominantly target excitatory neurons in the target area.

**Figure 6.**
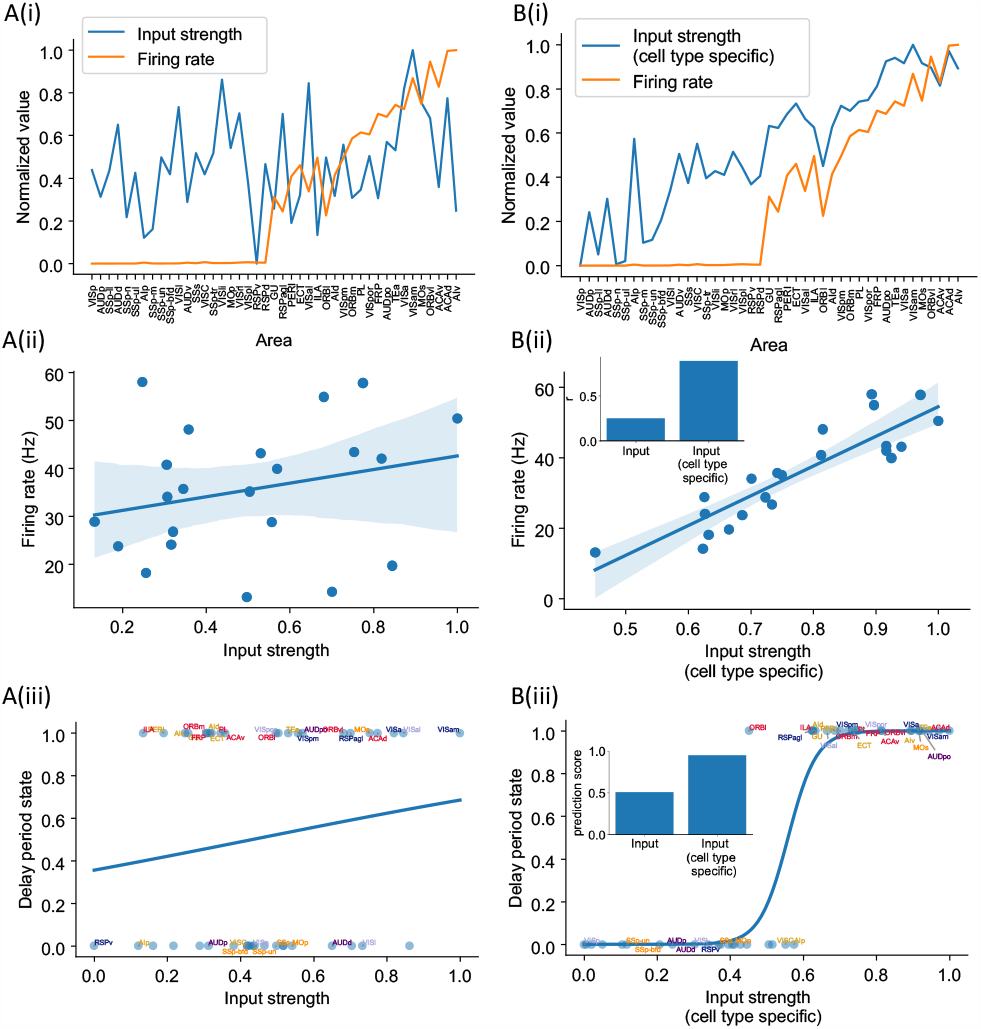
Cell type-specific connectivity measures are better at predicting firing rate pattern than nonspecific ones. (A(i)). Delay period firing rate (orange) and input strength for each cortical area. Input strength of each area is the sum of connectivity weights of incoming projections. Areas are plotted as a function of their hierarchical positions. Delay period firing rate and input strength are normalized for better comparison. (A(ii)). Input strength does not show significant correlation with delay period firing rate for areas showing persistent activity in the model (r = 0.25, p = 0.25). (A(iii)). Input strength cannot be used to predict whether an area shows persistent activity or not (prediction accuracy = 0.51). (B(i)). Delay period firing rate (orange) and cell type-specific input strength for each cortical area. Cell type-specific input strength considers how the long-rang connections target different cell types and is the sum of modulated connectivity weights of incoming projections. Same as (A(i)), areas are sorted according to their hierarchy and delay period firing rate and input strength are normalized for better comparison. (B(ii)). Cell type-specific input strength has a strong correlation with delay period firing rate of cortical areas showing persistent activity (r = 0.89, p < 0.05). Inset: Comparison of the correlation coefficient for raw input strength and cell type-specific input strength. (B(iii)). Cell type-specific input strength predicts whether an area shows persistent activity or not (prediction accuracy = 0.95). Inset: comparison of the prediction accuracy for raw input strength and cell type-specific input strength.

We found that cell type-specific graph measures accurately predict delay-period firing rates. The cell type-specific input strength of the early sensory areas is weaker than the raw input strength (Fig. 6B(i)). The firing rate across areas is positively correlated with cell type-specific input strength (Fig. 6B(ii)). Cell type-specific input strength also accurately predicts which areas show persistent activity (Fig. 6B(iii)). Similarly, we found that the cell type-specific eigenvector centrality, but not standard eigenvector centrality (Newman 2018), was a good predictor of delay period firing rates (Fig. 6 - supplement 2).

### A core subnetwork for persistent activity across the cortex

Many areas show persistent activity in our model. However, are all active areas equally important in maintaining persistent activity? When interpreting large-scale brain activity, we must distinguish different types of contribution to working memory. For instance, inactivation of an area like VISp impairs performance of a delay-dependent task because it is essential for a (visual) “input” to access working memory; on the other hand a “readout” area may display persistent activity only as a result of sustained inputs from other areas that form a “core”, which are causally important for maintaining a memory representation.

We propose four types of areas related to distributed working memory: input, core, readout, and nonessential (Fig. 7A). External stimuli first reach input areas, which then propagate activity to the core and non-essential areas. Core areas form recurrent loops and support distributed persistent activity across the network. By definition, disrupting any of the core areas would affect persistent activity globally. The readout areas also show persistent activity. Yet, inhibiting readout areas has little effect on persistent activity elsewhere in the network. We can assign the areas to the four classes based on three properties: a) the effect of inhibiting the area during stimulus presentation on delay activity in the rest of the network; b) the effect of inhibiting the area during the delay period on delay activity in the rest of the network; c) the delay activity of the area itself on trials without inhibition.

**Figure 7.**
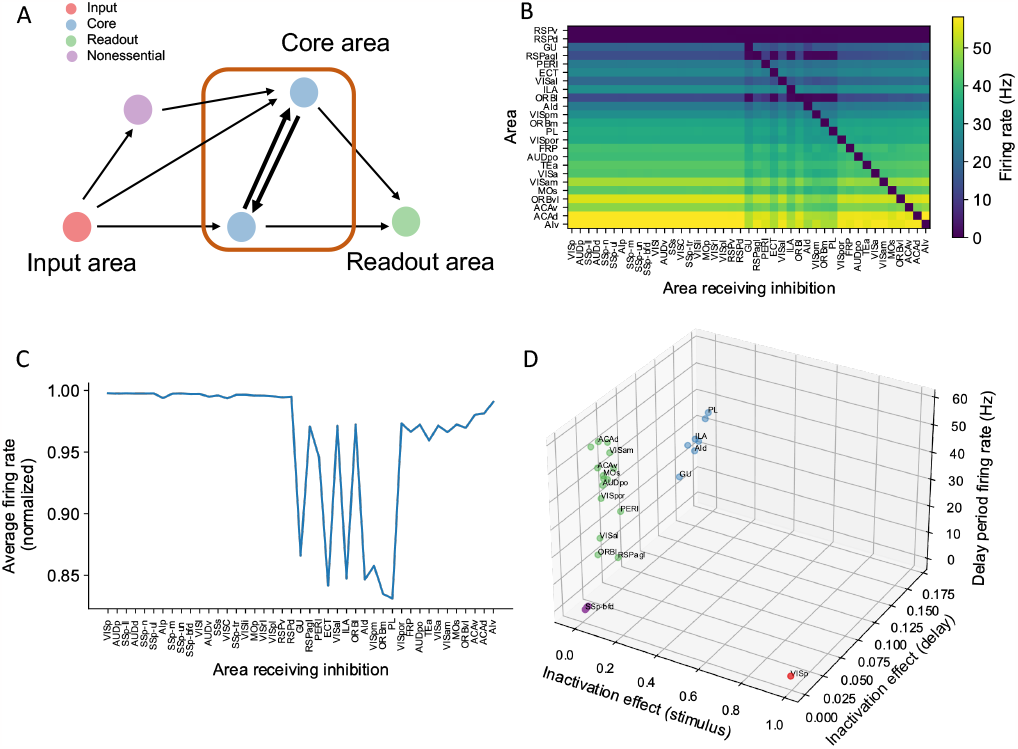
A core subnetwork generates persistent activity across the cortex. (A). We propose four different types of areas. Input areas (red) are responsible for coding and propagating external signals, which are then propagated through synaptic connections. Core areas (blue) form strong recurrent loops and generate persistent activity. Readout areas (green) inherit persistent activity from core areas. Nonessential areas (purple) may receive inputs and send outputs but they do not affect the generation of persistent activity. (B). Delay period firing rate for cortical areas engaged in working memory (Y axis) after inhibiting different cortical areas during the delay period (X axis). Areas in the X axis and Y axis are both sorted according to hierarchy. Firing rates of areas with small firing rate (<1Hz) are partially shown (only RSPv and RSPd are shown because their hierarchical positions are close to areas showing persistent activity). (C). The average firing rate for areas engaged in persistent activity under each inhibition simulation. The X axis shows which area is inhibited during the delay period, and the Y axis shows the average delay period activity for all areas showing persistent activity. Note that when calculating the average firing rate, the inactivated area was excluded in order to focus on the inhibition effect of one area on other areas. Average firing rates on the Y axis are normalized using the average firing in control (no inhibition) simulation. (D). Classification of 4 types of areas based on their delay period activity after stimulus- and delay-period inhibition (color denotes the type for area, as in A). The inhibition effect, due to either stimulus or delay period inhibition, is the change of average firing rate normalized by the average firing rate in the control condition. Areas with strong inhibition effect during stimulus period are classified as Input areas; areas with strong inhibition effect during delay period and strong delay period firing rate are classified as Core areas; areas with weak inhibition effect during delay period but strong delay period firing rate during control are classified as Readout areas; areas with weak inhibition effect during delay period and weak delay period firing rate during control are classified as Nonessential areas.

In search of a core working memory subnetwork in the mouse cortex, in model simulations we inactivated each area either during stimulus presentation or during the delay period, akin to optogenetic inactivation in mice experiments. The effect of inactivation was quantified by calculating the decrement in the firing rate compared to control trials for the areas that were not inhibited (Fig. 7B). The VISp showed a strong inhibition effect during the stimulus period, as expected for an Input area. We identified seven areas with a substantial inhibition effect during the delay period (Fig. 7C), which we identify as a core for working memory. Core areas are distributed across the cortex. They include frontal areas PL, ILA, medial part of the orbital area (ORBm), which are known to contribute to working memory (Liu et al. 2014; Bolkan et al. 2017). Other associative and sensory areas (AId, VISpm, ectorhinal area (ECT), gustatory area (GU)) are also in the core. Similarly, we used the above criteria to classify areas as Readout or Non-essential (Fig. 7D).

We have defined a core area for working memory maintenance as a cortical area that, first, exhibits persistent activity, and second, removal of this area (e.g., experimentally via a lesion or opto-inhibition) significantly affects persistent activity in other areas. It is possible, however, that effects on persistent activity at the network level only arise after lesioning two or more areas. Thus, we proceeded with inhibiting two, three, and four readout areas concurrently (Fig. 7-supplement 1A), as by definition, inhibiting any single readout area will not exhibit a strong inhibition effect.

We first inhibited pairs of readout areas and evaluated the effect of this manipulation at a network level. Specifically, for any given readout area A, we plotted the average firing rate of the network when A was inhibited as part of an inhibited pair (see description in Methods). After inhibiting a pair of readout areas, there was a decrement in the average firing rate of the network (Fig. 7 - supplement 1A). The decrement became more pronounced as more readout areas were inhibited, e.g., triplets and quadruplets, and when a combination of readout and core areas were inhibited pairwise (Fig. 7 - supplement 1B). This analysis demonstrates that readout areas also play a role in maintaining distributed persistent activity: we may define ’second-order core areas’ as those readout areas that have a strong inhibition effect only when inhibited concurrently with another area, while third-order and fourth-order core areas are analogously defined via triplet and quadruplet inhibition, respectively. We note that the effects of silencing pairs, triplets and quadruplets of readout areas remain smaller than those seen after silencing single core areas listed above. We also tested the effect of inhibiting all core areas during the delay period (Fig. 7 - supplement 1C). After inhibiting all core areas, some readout areas lost persistent firing. Moreover, there was a 48% decrement in the average firing rate compared with a 15% decrement for a single core area and a 3% decrement for a single readout area. Thus, the pattern of persistent activity is more sensitive to perturbations of core areas, which underscores the classification of some cortical areas into core vs readout.

### The core subnetwork can be identified by the presence of strong excitatory loops

Inhibition protocols across many areas are computationally costly. We sought a structural indicator that is easy to compute and is predictive of whether an area is engaged in working memory function. Such an indicator could also guide the interpretation of large-scale neural recordings in experimental studies. In the dynamical regime where individual cortical areas do not show persistent activity independently, distributed working memory patterns must be a result of long-range recurrent loops across areas. We thus introduced a quantitative measurement of the degree to which each area is involved in long-range recurrent loops (Fig. 8A).

**Figure 8.**
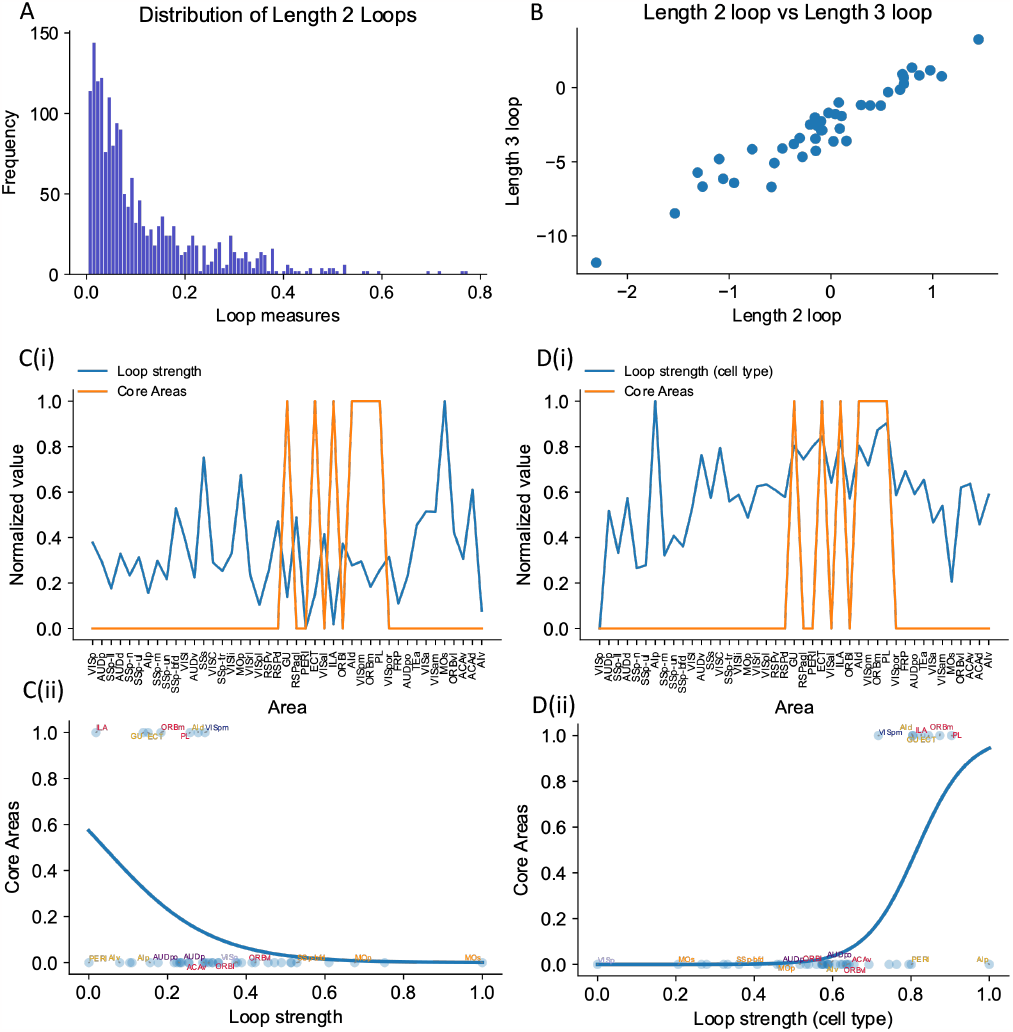
The core subnetwork can be identified structurally by the presence of strong excitatory loops. (A). Distribution of length-2 loops. X axis is the single loop strength of each loop (product of connectivity strengths within loop) and Y axis is their relative frequency. (B). Loop strengths of each area calculated using different length of loops (e.g., length 3 vs length 2) are highly correlated (r = 0.96, p < 0.05). (C(i)). Loop strength (blue) is plotted alongside Core Areas (orange), a binary variable that takes the value 1 if the area is a Core Area, 0 otherwise. Areas are sorted according to their hierarchy. The loop strength is normalized to a range of (0, 1) for better comparison. (C(ii)). A high loop strength value does not imply that an area is a core area. Blue curve shows the logistic regression curve fits to differentiate the core areas versus non core areas. (D(i)). Same as (C), but for cell type-specific loop strength. (D(ii)). A high cell type-specific loop strength predicts that an area is a core area (prediction accuracy = 0.93). Same as (C), but for cell type-specific loop strength.

The core subnetwork can be identified by the presence of strong loops between excitatory cells. Here we focus on length-2 loops (Fig. 8A); the strength of a loop is the product of two connection weights for a reciprocally connected pair of areas; and the loop strength measure of an area is the sum of the loop strengths of all length-2 loops that the area is part of. Results were similar for longer loops (Fig. 8B, also see Fig. 8 - supplement 1 for results of longer loops). The raw loop strength had no positive statistical relationship to the core working memory subnetwork (Fig. 8C(i), Fig. 8C(ii)). We then defined cell type-specific loop strength (see Methods). The cell type-specific loop strength is the loop strength calculated using connectivity multiplied by the cell type projection coefficient. The cell type-specific loop strength, but not the raw loop strength, predicts which area is a core area with high accuracy (Fig. 8D(i), Fig. 8D(ii), prediction accuracy = 0.93). This demonstrates that traditional connectivity measures are informative but not sufficient to explain dynamics during cognition in the mouse brain. Cell type-specific connectivity, and new metrics that account for such connectivity, are necessary to infer the role of brain areas in supporting large-scale brain dynamics during cognition.

To better demonstrate our cell type-specific connectivity measures, we have implemented two other measures for comparison: a) a loop-strength measure that adds a ‘sign’ without further modification, and b) a loop strength measure that takes hierarchical information - and not PV information-into account. These two graph-theoretic measures can be used to predict delay firing rate during a sensory working memory task, thus highlighting the importance of hierarchical information, which distinguishes excitatory from inhibitory feedback (Fig. 6 - supplement 3). On the other hand, the prediction of the core areas greatly depends on cell-type specificity: the sign-only and ‘no-PV’ mechanisms do not reliably predict whether an area is a core area or not, especially in the case of calculating with length 3 loops, demonstrating the importance of cell-type specific connectivity measures. (Fig. 8 - supplement 2).

### Multiple attractor states emerge from the mouse mesoscopic connectome and local recurrent interactions

Different tasks lead to dissociable patterns of internally sustained activity across the brain, described dynamically as distinct attractor states. Generally, attractor states may enable computations such as decision making and working memory (Wang 1999; Wang 2002; Mejias et al. 2016). Specifically, a given task may be characterized by a specific attractor landscape and thereby define different core areas for working memory, as introduced above. We developed a protocol to identify multiple attractor states, then analyzed the relationship between network properties and the attractor states (Fig. 9A-C). For different parameters, the number of attractors and the attractor patterns change. Two parameters are especially relevant here. These are the long-range connection strength (*μ*_*EE*_) and local excitatory connection strength (*g*_*E,self*_). These parameters affect the number of attractors in a model of the macaque cortex (Mejias and Wang 2022). Increasing the long range connection strength decreases the number of attractors (Fig. 9D). Stronger long-range connections implies that the coupling between areas is stronger. If areas are coupled with each other, the activity state of an area will be highly correlated to that of its neighbors. This leads to less variability and fewer attractors.

**Figure 9.**
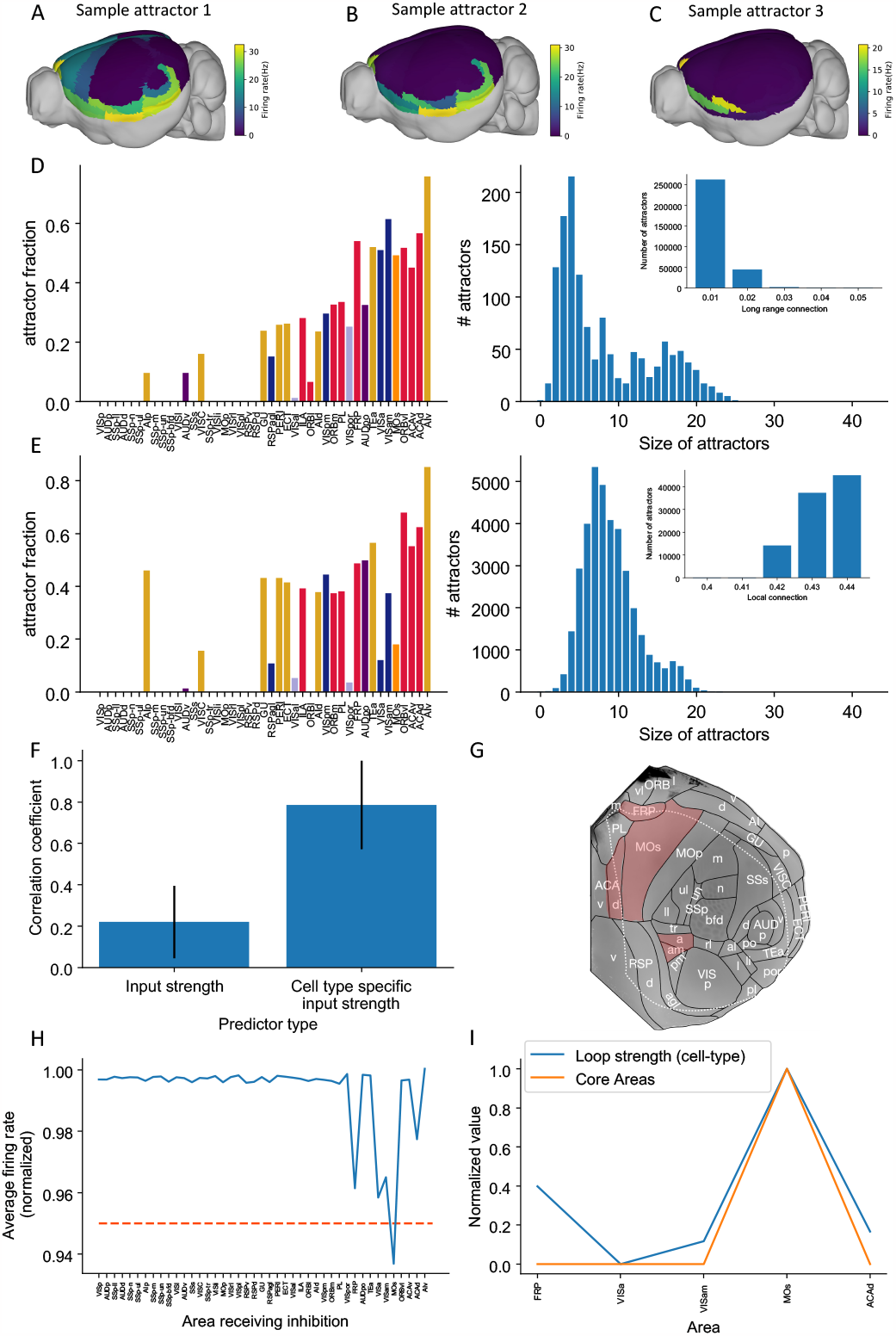
Multiple attractors coexist in the mouse working memory network. (A-C) Example attractor patterns with a fixed parameter set. Each attractor pattern can be reached via different external input patterns applied to the brain network. Delay activity is shown on a 3D brain surface. Color represents the firing rate of each area. (D-E) The distribution of attractor fractions (left) and number of attractors as a function of size (right) for different parameter combinations are shown. Attractor fraction of an area is the ratio between the number of attractors that include the area and the total number of identified attractors. In (D), local excitatory strengths are fixed (g_E,self_ = 0.44 nA) while long-range connection strengths vary in the range μ_EE_ = 0.01-0.05 nA. Left and right panels of (D) show one specific parameter μ_EE_ = 0.03 nA. Inset panel of (D) shows the number of attractors under different long-range connection strengths while g_E,self_ is fixed at 0.44 nA. In (E), long range connection strengths are fixed (μ_EE_ = 0.02 nA) while local excitatory strengths varies in the range g_E,self_ = 0.4-0.44 nA. Left and right panels of (E) show one specific parameter g_E,self_ = 0.43 nA. Inset panel of (E) shows the number of attractors under different local excitatory strengths, while μ_EE_ is fixed at 0.02 nA. (F). Prediction of the delay period firing rate using input strength and cell type-specific input strength for each attractor state identified under μ_EE_ = 0.04 nA and g_E,self_ = 0.44 nA. 143 distinct attractors were identified and the average correlation coefficient using cell type-specific input strength is better than that using input strength. (G). A example attractor state identified under the parameter regime μ_EE_ = 0.03 nA and g_E,self_ = 0.44 nA. The 5 areas with persistent activity are shown in red. (H). Effect of single area inhibition analysis for the attractor state in (G). For a regime where 5 areas exhibit persistent activity during the delay period, inactivation of the premotor area MOs yields a strong inhibition effect (<0.95 orange dashed line) and is therefore a Core area for the attractor state in (G). (I). Cell type-specific loop strength (blue) is plotted alongside Core areas (orange) for the attractor state in (G). Only 5 areas with persistent activity are used to calculate the loop strength. Loop strength is normalized to be within the range of 0 and 1. High cell type-specific loop measures predict that an area is a Core area (prediction accuracy is 100% correct). The number of areas is limited, so prediction accuracy is very high.

To quantify how the patterns of attractors change for different parameters, two quantities are introduced. The *attractor fraction* is the fraction of all detected attractor states to which an area belongs. An area “belongs” to an attractor state if it is in a high activity state in that attractor. The *attractor size* is defined by the number of areas belonging to that attractor. As we increased the long-range connection strength, the attractor size distribution became bimodal. The first mode corresponded to large attractors, with many areas. The second mode corresponded to small attractors, with few areas (Fig. 9D).

When the local excitatory strength is increased, the number of attractors increased as well (Fig. 9E). In this regime some areas are endowed with sufficient local reverberation to sustain persistent activity even when decoupled from the rest of the system, therefore the importance of long-range coupling is diminished and a greater variety of attractor states is enabled. This can be understood by a simple example of two areas 1 and 2, each capable of two stimulus-selective persistent activity states; even without coupling there are 2 *×* 2 = 4 attractor states with elevated firing. Thus, local and long-range connection strength have opposite effects on the number of attractors.

The cell type-specific input strength predicted firing rates across many attractors. In an example parameter regime (*μ*_*EE*_ = 0.04 nA and *g*_*E,self*_ = 0.44 nA), we identified 143 attractors. We correlated the input strength and cell type-specific input strength with the many attractor firing rates (Fig. 9F). The raw input strength is weakly correlated with activity patterns. The cell type-specific input strength is strongly correlated with activity across attractors. This shows that the cell type-specific connectivity measures are better at predicting the firing rates in many scenarios. These results further prove the importance of having cell type-specific connectivity for modeling brain dynamics.

Different attractor states rely on distinct subsets of core areas. In one example attractor, we found 5 areas that show persistent activity: VISa, VISam, FRP, MOs and ACAd (Fig. 9G) (parameter regime, *μ*_*EE*_ = 0.03 nA and *g*_*E,self*_ = 0.44 nA). We repeated the previous inhibition analysis to identify core areas for this attractor state. Inhibiting one area, MOs, during the delay had the strongest effect on delay activity in the other parts of the attractor (Fig. 9H). MOs also showed strong persistent activity during delay period. This is consistent with its role in short-term memory and planning (Li et al. 2015; Inagaki et al. 2019). According to our definition, MOs is a core area for this attractor. To calculate a loop strength that was specific to this attractor, we only examined connections between these five areas. The cell type-specific loop strength was strongest in area MOs (Fig. 9I). Thus, we can identify likely core areas for individual attractor states from cell type-specific structural measures. This also demonstrates that different attractor states can be supported by distinct core areas.

## Discussion

We developed a connectome-based dynamical model of the mouse brain. The model was capable of internally maintaining sensory information across many brain areas in distributed activity in the absence of any input. To our knowledge this is the first biologically-based model of the entire mouse cortex and the thalamocortical system that supports a cognitive function, in this case working memory. Together with our recent work (Mejias and Wang 2022; Froudist-Walsh et al. 2021; Froudist-Walsh et al. 2023), it provides an important reference point to study the differences between mice and monkeys.

Our main findings are threefold. First, the mnemonic activity pattern is shaped by the differing densities of PV interneurons across cortical areas. Areas with a high PV cell fraction encoded information only transiently. Those with low PV cell fraction sustained activity for longer periods. Thus, the gradient of PV cells (Kim et al. 2017) has a definitive role in separating rapid information processing in sensory areas from sustained mnemonic information representation in associative areas of the mouse cortex. This is consistent with the view that each local area operates in the “inhibition-stabilizing regime” where recurrent excitation alone would lead to instability but the local network is stabilized by feedback inhibition, which may arise from long-range excitatory inputs to inhibitory neurons. This consistent with the regime of the primary visual cortex (R. J. Douglas et al. 1995; Murphy and Miller 2009). Second, we deliberately considered two different dynamical regimes: when local recurrent excitation is not sufficient to sustain persistent activity and when it is. In the former case, distributed working memory must emerge from long-range interactions between parcellated areas. Thereby the concept of synaptic reverberation (Lorente de Nó 1933; P. S. Goldman-Rakic 1995; Wang 2001; Wang 2021) is extended to the large-scale global brain. Note that currently it is unclear whether persistent neural firing observed in a delay dependent task is generated locally or depends on long-distance reverberation among multiple brain regions. Our work made the distinction explicit and offers specific predictions to be tested experimentally. Third, presently available connectomic data are not sufficient to account for neural dynamics and distributed cognition, and we propose cell type-specific connectomic measures that are shown to predict the observed distributed working memory representations. Our model underscores that, although connectome databases are an invaluable resource for basic neuroscience, they should be supplemented with cell-type-specific information.

We found that recurrent loops within the cortex and the thalamocortical network aided in sustaining activity throughout the delay period (Guo et al. 2017; Schmitt et al. 2017). The presence of thalamocortical connections had a similar effect on the model as cortico-cortical projections, with the distinct contributions of the thalamus to large-scale dynamics still to be uncovered (Shine et al. 2018; Jaramillo et al. 2019). The specific pattern of cortico-cortical connections was also critical to working memory. However, standard graph theory measures based on the connectome were unable to predict the pattern of working memory activity. By focusing on cell type-specific interactions between areas, we were able to reveal a core of cortical areas. The core is connected by excitatory loops, and is responsible for generating a widely distributed pattern of sustained activity. This clarifies the synergistic roles of the connectome and gradients of local circuit properties in producing a distributed cognitive function.

Previous large-scale models of the human and macaque cortex have replicated functional connectivity (Deco et al. 2014; Demirtaş et al. 2019; Honey et al. 2007; Schmidt et al. 2018; Shine et al. 2018; Cabral et al. 2011; Wang et al. 2019) and propagation of information along the cortical hierarchy (Chaudhuri et al. 2015; Joglekar et al. 2018; Diesmann et al. 1999). More recently, large-scale neural circuit models have been developed specifically to reproduce neural activity during cognitive tasks (Mejias and Wang 2022; Froudist-Walsh et al. 2021; Klatzmann et al. 2022). These models consider the fact that in the macaque cortex, the density of spines on pyramidal cells increases along the cortical hierarchy (Elston and Rosa 1998; Elston 2007; Chaudhuri et al. 2015). In a large-scale model of the macaque cortex (Chaudhuri et al. 2015), it was shown that this ‘excitatory gradient’ (Wang 2020) is correlated with the distribution of intrinsic timescales in the cortex (Murray et al. 2014) and is consistent with spatially distributed working memory patterns (Mejias and Wang 2022; Froudist-Walsh et al. 2021). Such excitatory gradients based on spine count are less pronounced, and may be entirely absent in the rodent cortex (Ballesteros-Yáñez et al. 2010; Gilman et al. 2017). However, there are gradients of synaptic inhibition in the mouse cortex (Kim et al. 2017; Wang 2021). Kim et al., showed that the ratio of SST+ neurons to PV+ neurons is low for early sensory areas and motor areas, while it is high in association areas such as the frontal cortex. We have used this gradient of inhibition in our model to show that spatially distributed persistent-activity patterns in the mouse cortex do not require gradients of recurrent excitation. In our model, the PV gradient and CIB may be particularly important to maintain the stability of an otherwise highly excitable cortical area. Along these lines, we predict that local recurrency in the mouse early sensory areas is higher than in the primate. Consistent with this claim, both the spine density and the number of excitatory and inhibitory synapses in layer 2/3 pyramidal neurons in area V1 are higher in mouse compared to macaque (Fig. 5A in Gilman et al. 2017, Fig. 1A in Wildenberg et al. 2021).

Other anatomical properties at the area and single cell level may be informative of the differences in computational and/or cognitive abilities between rodents and macaques. In the language of network theory, the macaque cortex is a densely connected graph at an inter-area level, with the connectivity spanning five orders of magnitude (Markov et al. 2014a), which is more than what is expected for small-world networks (Bassett and Bullmore 2017). Critically, the mouse ‘connectome’ (e.g., (Oh et al. 2014; Harris et al. 2019; Knox et al. 2018) has even denser area-to-area connections. In the visual cortex, individual neurons target more cortical areas in the mouse (Siu et al. 2021) and they have more inhibitory and excitatory synapses (Wildenberg et al. 2021). Thus, connectivity in the mouse is denser at both the area and single-cell levels (at least for primary visual cortex). We propose that there is a greater functional specialization in the primate cortex which is afforded by the sparser and more targeted patterns of connectivity at the single-cell and area levels. Other differences to explore in future computational models include the ratio of NMDA to AMPA-mediated synaptic currents, which is approximately constant in the mouse cortex (Myme et al. 2003) but varies along the cortical hierarchy in primates (Yang et al. 2018; Klatzmann et al. 2022), as well as hierarchy, which is defined based on feedforward and feedback projections in the mouse (Harris et al. 2019) and primate (Markov et al. 2014a).

We found that traditional graph theory metrics of connectivity were unable to predict the working memory activity in the mouse brain. This may be due to the almost fully connected pattern of interareal connectivity in the mouse cortex (Gămănuţ et al. 2018). This implies that, qualitatively, all areas have a similar set of cortical connections. In our model, we allowed the cell type target of interareal connections to change according to the relative position of the areas along the cortical hierarchy. Specifically, feedforward connections had a greater net excitatory effect than feedback connections, a hypothesis which we refer to as CIB. This preferential targeting of feedback projections serves to stabilize the otherwise excitable activity of sensory areas (Mejias and Wang 2022), and is consistent with recent experiments that report long-range recruitment of GABAergic neurons in early sensory areas (Campagnola et al. 2022; Shen et al. 2022; Naskar et al. 2021). Our model predicts that if there is a weak correlation between PV cell density and delay firing rate across cortical areas, then the CIB mechanism is at play. Moreover, the model results suggest that CIB is particularly important in the regime where local connections are not sufficient to sustain spatially-patterned persistent activity. We also showed that there are parameter regimes where CIB becomes less important, provided there is a gradient of synaptic inhibition as in the mouse cortex ((Kim et al. 2017), but see (Nigro et al. 2022)). Notably, the model’s resilience to parameter variations in inhibitory connection strengths is significantly enhanced when both the PV gradient and CIB are present. Given that working memory is a fundamental cognitive function observed across many individual brains with anatomical differences, the inclusion of multiple inhibitory mechanisms that allow for connectivity variations might confer evolutionary advantages. Although there is some evidence for similar inhibitory gradients in humans (Burt et al. 2018) and macaque (Torres-Gomez et al. 2020), the computational consequences of differences across species remain to be established.

To conclude, the manner in which long-range recurrent interactions affect neural dynamics depends not only on the existence of excitatory projections per se, but also on the target neurons’ cell type. Thus, for some cortical areas afferent long-range excitatory connections promote working memory-related activity while for some others, e.g., early sensory areas, it does not. Moreover, the existence of long-range interactions is consistent with potentially distinct dynamical regimes. For example, in one regime some areas exhibit independent persistent activity, i.e., local recurrent interactions are sufficient to sustain a memory state for these areas, while others do not. In this regime CIB is not required for the existence of distributed persistent activity patterns. In another regime, none of the areas can sustain a memory state without receiving long-range input. These two regimes are functionally distinct in terms of their robustness to perturbation as well as in the number of attractors that they can sustain. These regimes may be identified via perturbation analysis in future experimental and theoretical work.

By introducing cell type-specific graph theory metrics, we were able to predict the pattern and strength of delay period activity with high accuracy. Moreover, we demonstrated how cell type-specific graph-theory measures can accurately identify the core subnetwork, which can also be identified independently using a simulated large-scale optogenetic experiment. We found a core subnetwork of areas that, when inhibited, caused a substantial drop in activity in the remaining cortical areas. This core working memory subnetwork included frontal cortical areas with well documented patterns of sustained activity during working memory tasks, such as prelimbic (PL), infralimbic (ILA) and medial orbitofrontal cortex (ORBm) (Schmitt et al. 2017; Liu et al. 2014; Wu et al. 2020). However, the core subnetwork for the visual working memory task we assessed was distributed across the cortex. It also included temporal and higher visual areas, suggesting that long-range recurrent connections between the frontal cortex and temporal and visual areas are responsible for generating persistent activity and maintaining visual information in working memory in the mouse.

Some of the areas that were identified as core areas in our model have been widely studied in other tasks. For example, the gustatory area exhibits delay-period preparatory activity in a taste-guided decision-making task and inhibition of this area during the delay period impairs behavior (Vincis et al. 2020).

The core visual working memory subnetwork generates activity that is then inherited by many readout areas, which also exhibit persistent activity. However, inhibiting readout areas only mildly affects the activity of other areas (Fig. 7 and Fig. 7 - supplement 1). The readout areas in our model were a mixture of higher visual areas, associative areas and premotor areas of cortex. We also concluded that MOs is a readout area and not a core area. This finding may be surprising considering previous studies that have shown this area to be crucial for short-term memory maintenance, planning, and movement execution during a memory-guided response task (Guo et al. 2017; Guo et al. 2014; Inagaki et al. 2019; Li et al. 2015; Wu et al. 2020; Voitov and Mrsic-Flogel 2022). This task has shown to engage, not only ALM, but a distributed subcortical-cortical network that includes the thalamus, basal ganglia and cerebellum (Svoboda and Li 2018). We note that in the version of the memory-guided response task studied by Svoboda and others, short-term memory is conflated with movement preparation. In our task, we proposed to study the maintenance of sensory information independent of any movement preparation as in delayed match-to-sample tasks and variations thereof. It is for this behavioral context that we found that MOs is not a core visual working memory area. We emphasize that readout areas are not less important than core areas as readout areas can use the stored information for further computations and thus some readout areas are expected to be strongly coupled to behavior. Indeed, there is evidence for a differential engagement of cortical networks depending on the task design (Jonikaitis et al. 2023) and on effectors (Kubanek and Snyder 2015). If ALM is indeed a readout area for sensory working memory tasks, (e.g., (Schmitt et al. 2017)), then the following prediction arises. Inhibiting ALM should have a relatively small effect on sustained activity in core areas (such as PL) during the delay period. In contrast, inhibiting PL and other core areas may disrupt sustained activity in ALM. Even if ALM is not part of the core for sensory working memory, it could form part of the core for motor preparation tasks (Fig. 9G). We found a high cell-type-specific loop strength for area ALM, like that in core areas, which supports this possibility (Fig. 9I). Furthermore, we found some attractor states for which the MOs was classified as a core area, that do not contain area PL. This result is supported by a recent study that found no behavioral effect after PL inhibition in a motor planning task (Wang et al. 2021). Therefore, the core subnetwork required for generating persistent activity is likely task-dependent. Future modeling work may help elucidate the biological mechanisms responsible for switching between attractor landscapes for different tasks.

Neuroscientists are now observing task-related neural activity at single-cell resolution across much of the brain (Stringer et al. 2019; Steinmetz et al. 2019). This makes it important to identify ways to distinguish the core areas for a function from those that display activity that serves other purposes. We show that a large-scale inhibition protocol can identify the core subnetwork for a particular task. We further show how this core can be predicted based on the interareal loops that target excitatory neurons. Were such a cell type-specific interareal connectivity dataset available, it may help interpretation of large-scale recording experiments. This could also focus circuit manipulation on regions most likely to cause an effect on the larger network activity and behavior. Our approach identifies the brain areas that work together to support working memory. It also identifies those that benefit from such activity to serve other purposes. Our simulation and theoretical approach is therefore ideally suited to understand the large-scale anatomy, recording and manipulation experiments which are at the forefront of modern systems neuroscience.

Neuroscience has rapidly moved into a new era of investigating large-scale brain circuits. Technological advances have enabled the measurement of connections, cell types and neural activity across the mouse brain. We developed a model of the mouse brain and theory of working memory that is suitable for the large-scale era. Previous reports have emphasized the importance of gradients of dendritic spine expression and interareal connections in sculpting task activity in the primate brain (Mejias and Wang 2022; Froudist-Walsh et al. 2021). Although these anatomical properties from the primate cortex are missing in the mouse brain (Gămănuţ et al. 2018; Gilman et al. 2017), other properties such as interneuron density (Kim et al. 2017) may contribute to areal specialization. Indeed, our model clarifies how gradients of interneurons and cell type-specific interactions define large-scale activity patterns in the mouse brain during working memory, which enables sensory and associative areas to have complementary contributions. Future versions of the large-scale model may consider different interneuron types to understand their contributions to activity patterns in the cortex (Kim et al. 2017; Meng et al. 2023; Froudist-Walsh et al. 2021; Wang et al. 2004; Tremblay et al. 2016; Nigro et al. 2022), the role of interhemispheric projections in providing robustness for short-term memory encoding (Li et al. 2016a), and the inclusions of populations with tuning to various stimulus features and/or task parameters that would allow for switching across tasks (Yang et al. 2019). Importantly, these large-scale models may be used to study other important cognitive computations beyond working memory, including learning and decision making (Abbott et al. 2017; Abbott et al. 2020).

## Acknowledgements

We thank Daniel P. Bliss and Ulises Pereira for support with analysis tools at the beginning of the project, and members of the Wang Lab at New York University for discussions related to the project. This work was funded by US National Institutes of Health (NIH) grant R01MH062349, Office of Naval Research (ONR) grant N00014, National Science Foundation (NSF) NeuroNex grant 2015276, Simons Foundation grant 543057SPI, and NIH grant U19NS123714 to JJ and X-JW and a Bristol Neuroscience of Mental Health award and UK Research and Innovation (UKRI) Biotechnology and Biological Sciences Research Council (BBSRC) grant BB/X013243/1 to S.F.W.

## Author contributions

XJW: designed the research, worked with the other authors throughout the project and co-wrote the paper; XYD: designed the model, carried out all the computer simulations and analysis of simulation data, and co-wrote the paper; SFW provided initial simulation code, analyzed simulation data, supervised research, and co-wrote the paper; JJ analyzed simulation data, supervised research, and co-wrote the paper; JJJ: contributed analytic tools.

## Methods

### Anterograde tracing, connectivity data

We used the mouse connectivity map from Allen institute (Oh et al. 2014) to constrain our large-scale circuit model of the mouse brain. The Allen Institute measured the connectivity among cortical and subcortical areas using an anterograde tracing method. In short, they injected virus and expressed fluorescent protein in source areas and performed fluorescent imaging in target areas to measure the strength of projections from source areas. Unlike retrograde tracing methods used in other studies (Markov et al. 2014b), the connectivity strength measured using this method does not need to be normalized by the total input or output strength. This means that connectivity strength between any two areas is comparable.

The entries of the connectivity matrix from the Allen Institute can be interpreted as proportional to the total number of axonal fibers projecting from unit volume in one area to unit volume in another area. Before incorporating the connectivity into our model, we normalized the data as follows. In each area, we model the dynamics of an “average” neuron, assuming that the neuron receive inputs from all connected areas. Thus, we multiplied the connectivity matrix by the volume *V ol*_*j*_ of source area *j* and divided by the average neuron density *d*_*i*_ in target area *i*:

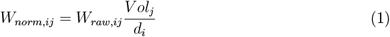

where *W*_*raw,ij*_ is the raw, i.e., original, connection strength from unit volume in source area *j* to unit volume in target area *i, V ol*_*j*_ is the volume of source area *j* (Wang et al. 2020), and *d*_*i*_ is the neuron density in source area *i* (Erö et al. 2018). *W*_*norm,ij*_ is the matrix that we use to set the long rang connectivity in our circuit model. We can define the cortico-thalamic connectivity *W*_*ct,norm,ij*_ and thalamo-cortical connectivity *W*_*tc,norm,ij*_ in a similar manner, except that we didn’t apply the normalization to thalamic connectivity due to not having enough neuron density data.

### Interneuron density along the cortex

Kim and colleagues measured the density of typical interneuron types in the brain (Kim et al. 2017). They expressed fluorescent proteins in genetically labeled interneurons and counted the number of interneurons using fluorescent imaging. We took advantage of these interneuron density data and specifically used the PV cell fraction to set local and long-range inhibitory weights.

The PV cell density of all layers is first divided by the total neuron density *d*_*i*_ in the area *i*, to give the PV cell fraction *PV*_*raw,i*_, which better reflects the expected amount of synaptic inhibition mediated by PV neurons. The PV cell fraction is then normalized across the whole cortex.

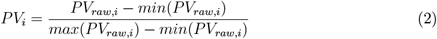

*P V*_*raw,i*_ is the PV cell fraction in area *i*, and *P V*_*i*_ is the normalized value of *P V*_*raw*_, which will be used in subsequent modeling.

### Hierarchy in the cortex

The concept of hierarchy is important for understanding the cortex. Hierarchy can be defined based on mapping corticocortical long range connections onto feedforward or feedback connections (Felleman and Essen 1991; Markov et al. 2014a; Harris et al. 2019). Harris and colleagues measured the corticocortical projections and target areas in a series of systematic experiments in mice (Harris et al. 2019). Projection patterns were clustered into multiple groups and the label “feedforward” or “feedback” was assigned to each group. Feedforward and feedback projections were then used to determine relative hierarchy between areas. For example, if the projections from area A to area B are mostly feedforward, then area B has a higher hierarchy than area A. This optimization process leads to a quantification of the relative hierarchy of cortical areas *h*_*raw,i*_. We defined the normalized hierarchy value *h*_*i*_ as

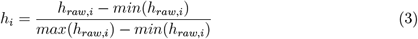

where *h*_*raw,i*_ is the raw, i.e., original hierarchical ordering from (Harris et al. 2019). Due to data acquisition issues, 6 areas did not have a hierarchy value assigned to them (SSp-un, AUDv, GU, VISC, ECT, PERI) (Harris et al. 2019). We estimated hierarchy through a weighted sum of the hierarchy value of 37 known areas, while the weight is determined through the connectivity strength. The parameters *α*_*h*_ and *β*_*h*_ are selected so that *h*_*i,estimate*_ are close to *h*_*i*_ for areas with known hierarchy.

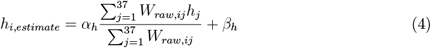

For the thalamocortical model, we also used the hierarchy value for thalamic areas (Harris et al. 2019). The hierarchy of thalamic areas are comparable to cortical areas, so in order to use it in the model, we also normalized them.

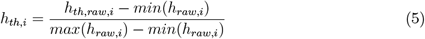

To estimate the hierarchy value of thalamic areas with missing values, we used the known hierarchy value of the thalamic area next to the missing one as a replacement.

### Description of the local circuit

Our large-scale circuit model includes 43 cortical areas. Each area includes two excitatory populations, labeled A and B, and one inhibitory population, C. The two excitatory populations are selective to different stimuli. The synaptic dynamics between populations are based on previous firing rate models of working memory (Wang 1999; Wong and Wang 2006). The equations that define the dynamics of the synaptic variables are

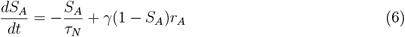

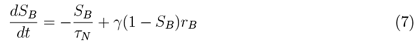

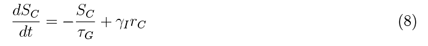

where *S*_*A*_ and *S*_*B*_ are the NMDA synaptic variables of excitatory populations A and B, while *S*_*C*_ is the GABA synaptic variable of the inhibitory population C. *r*_*A*_, *r*_*B*_ and *r*_*C*_ are the firing rates of populations A, B and C, respectively. *τ*_*N*_ and *τ*_*G*_ are the time constants of NMDA and GABA synaptic conductances. *γ* and *γ*_*I*_ are the parameters used to scale the contribution of presynaptic firing rates. The total currents received *I*_*i*_ (*i* = *A, B, C*) are given by

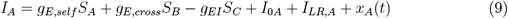

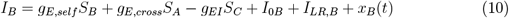

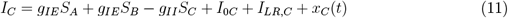

In these equations, *g*_*E,self*_, *g*_*E,cross*_ denote the connection strength between excitatory neurons with same or different selectivity, respectively. These connection strengths are the same for different areas, since there is no significant gradient for excitatory strength in mice. *g*_*IE*_ are the connection strengths from excitatory to inhibitory neurons, while *g*_*EI*_, and *g*_*II*_ are connection strengths from inhibitory to excitatory neurons and from inhibitory to inhibitory neurons, respectively. These connections will be scaled by PV cell fraction *PV*_*i*_ in the corresponding area. We will discuss the details in the next section. *I*_0*i*_ (*i* = *A, B, C*) are constant background currents to each population. *I*_*LR,i*_ (*i* = *A, B, C*) are the long range (LR) currents received by each population. The term *x*_*i*_(*t*) where *i* = *A, B, C* represents noisy contributions from neurons external to the network. It is modeled as an Ornstein-Uhlenbeck process:

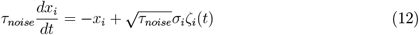

where *ζ*_*i*_(*t*) is Gaussian white noise, *τ*_*noise*_ describes the time constant of external AMPA synapses and *σ*_*i*_ sets the strength of the noise for each population. *σ*_*A*_ = *σ*_*B*_ = 5*pA* while *σ*_*C*_ = 0*pA*.

The steady state firing rate of each population is calculated based on a transfer function *ϕ*_*i*_(*I*) of input current received by each population *I*_*i*_ (*i* = *A, B, C*) given by

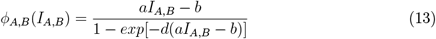

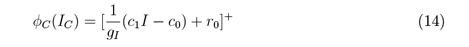

Note that the transfer functions *ϕ*_*i*_(*t*) are the same for two excitatory populations. *x*^+^ denotes the positive part of the function *x*. The firing rate of each population follows equations:

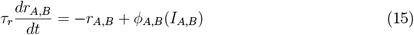

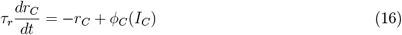

### Interneuron gradient and local connections

We scaled local interneuron connectivity with the interneuron density that was obtained using fluorescent labeling (Kim et al. 2017). Specifically, local I-I connections and local I-E connections are scaled by the interneuron density by setting the connection strength *g*_*k,i*_(*k* = *EI, II*) as a linear function of PV cell fraction *P V*_*i*_ in area *i*.

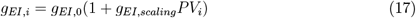

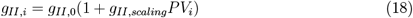

where *g*_*k*,0_ (*k* = *EI, II*) is the base value of I to E connections and *g*_*k,scaling*_ (*k* = *EI, II*) is the scaling factor of PV value. *g*_*k*,0_ also accounts for the inhibition of other cell types not explicitly considered in this study.

### Hierarchy and long range connections

Long range (LR) connections between areas are scaled by connectivity data from the Allen Institute (Oh et al. 2014). We consider long-range connections that arise from excitatory neurons because most long-range connections in the cortex correspond to excitatory connections (Petreanu et al. 2009). Long-range connections will target excitatory populations in other brain areas with the same selectivity (Zandvakili and Kohn 2015) and will also target inhibitory neurons. These long-range connections are given by the following equations:

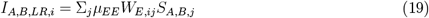

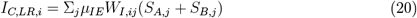

where *W*_*E*_ is the normalized long-range connectivity to excitatory neurons, and *W*_*I*_ is the normalized long-range connectivity to inhibitory neurons. *μ*_*EE*_ and *μ*_*IE*_ are coefficients scaling the long-range E to E and E to I connection strengths, respectively.

Here, we assume that the long-range connections will be scaled by a coefficient that is based on the hierarchy of source and target area. To quantify the difference between long-range feedforward and feedback projections, we introduce *m*_*ij*_ to measure the “feedforwardness” of projections between two areas. According to our assumption of counterstream inhibitory bias (CIB), long-range connections to inhibitory neurons are stronger for feedback connections and weaker for feedforward connections, while the opposite holds for long range connections to excitatory neurons. Following this hypothesis, we define *m*_*ij*_ as a sigmoid function of the difference between the hierarchy value of source and target areas. For feedforward projections, *m*_*ij*_ *>* 0.5; for feedback projections, *m*_*ij*_ *<* 0.5. Excitatory and inhibitory long-range connection strengths are implemented by multiplying the long-range connectivity strength *W*_*ij*_ by *m*_*ij*_ and (1 *− m*_*ij*_), respectively:

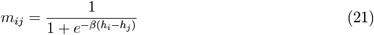

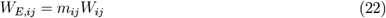

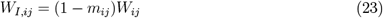

with

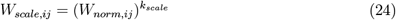

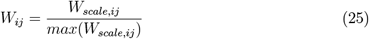

The connectivity *W*_*norm,ij*_ is then rescaled to translate the broad range of connectivity values (over five orders of magnitude) to a range more suitable for our firing rate models. *k*_*scale*_ is the coefficient used for this scaling. *k*_*scale*_ *<* 1 effectively makes the range much smaller than the original normalized connectivity *W*_*norm,ij*_. After that, the scaled connectivity *W*_*scale,ij*_ is then normalized so that the maximum value is fixed at 1.

### Simulations of replacing the PV gradient and CIB

In order to demonstrate the importance of PV gradient and CIB, we replace the PV gradient value/CIB with the average value accordingly in the simulation. Specifically, we replace PV gradient with the average PV cell fraction.

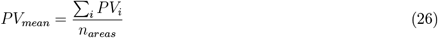

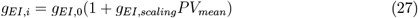

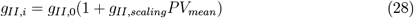

We also replace CIB with its average value 0.5, which means there is no bias to inhibitory cells for all long range connections.

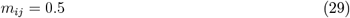

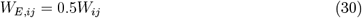

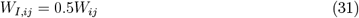

For the simulations of varying the local inhibitory connection strengths, we specifically change the value of *g*_*EI*,0_ and *g*_*EI,scaling*_ for *g*_*EI,i*_. For each combination of parameters of *g*_*EI*,0_ and *g*_*EI,scaling*_, simulations are performed for default parameters (no changes to PV gradient or CIB), PV average (PV gradient is replaced by average value) and CIB average (CIB is replaced by average value). The average firing rate of all areas and number of areas showing persistent activity are quantified for each parameter combination.

In other simulations, we varied the parameters *g*_*EI*,0_ and *g*_*EI,scaling*_ with long range connections *μ*_*EE*_ and *μ*_*IE*_ set to be 0. This enabled us to discover the range of parameter values for which individual areas were capable of maintaining persistent activity without input from other areas. In practice, the only key parameter that determines this behaviour is the smallest inhibitory connection strength of any area, *g*_*EI,i*_ = *g*_*EI*,0_.

### Simulations and theoretical calculation of the baseline stability of the network

In the simulation focusing on the stability of the baseline state of the network, there was no external input provided to any of the areas apart from noise (Eq. 12). The steady firing rate of each area after 10 s is recorded as a measure of the baseline stability.

We tested the baseline stability on five different scenarios (Fig. 4A-B) : In (1) and (2) we set the long-range connections *μ*_*EE*_ and *μ*_*IE*_ to zero since we focus on the local network. In (2), we also set the local inhibitory connections *g*_*EI*,0_ to zero. In (3) - (5) the long-range connections are intact. In (4), we set the long-range connection to inhibitory neurons *μ*_*IE*_ to zero. In (5), we set the local inhibitory connections *g*_*EI*,0_ to zero.

We analytically calculated the stability of baseline state for a local circuit when the long range connections *μ*_*EE*_ and *μ*_*IE*_ are set to zero, which means *I*_*LR,A*_, *I*_*LR,B*_, *I*_*LR,C*_ are zero in Eqs 9-11. In Eqs 15 and 16, we assume that *r*_*A*_, *r*_*B*_ and *r*_*C*_ reach their steady states instantaneously, since its time constant *τ*_*r*_ is much smaller than the time constant of NMDA synaptic variable *τ*_*N*_ in Eqs 6 and 7. Thus, we can express the firing rate *r*_*A*_, *r*_*B*_ and *r*_*C*_ as functions of synaptic variables *S*_*A*_, *S*_*B*_ and *s*_*C*_ (Eqs. 13-16):

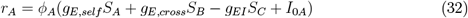

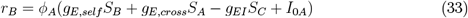

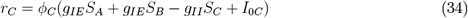

where *ϕ*_*A*_ and *ϕ*_*C*_ have the same form as Eqs 13 and 14.

Then we can insert Eqs 32-34 into Eqs 6-8 to obtain a differential equation for *S*_*A*_, *S*_*B*_ and *S*_*C*_.

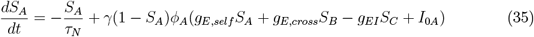

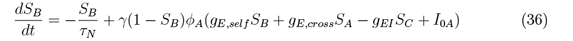

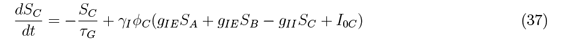

The steady state of *S*_*A*_, *S*_*B*_ and *S*_*C*_ can be solved numerically by setting the left side of the above equations to be zero. We denote the right side of the equations as *FA, FB* and *FC*. Then we can calculate the Jacobian matrix and its eigenvalues.

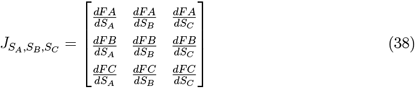

If the real part of all the eigenvalues are negative, then that means the baseline state is stable. The eigenvalues of the scenario (1) are -10.4, -12.5 and -229.8, while those of scenario (2), where local inhibitory connections *g*_*EI*,0_ are zero, are -7.4, -7.9, -232.3. These results coincide with the simulation results of Fig. 4A.

We also considered an alternative parameter regime, where the local excitatory connections *g*_*E,self*_ is set to a higher level *g*_*E,self*_ = 0.6*nA*. The local inhibitory connections strength *g*_*EI*,0_ is also set to a higher level *g*_*EI*,0_ = 0.5*nA* to balance the increased excitatory connections. Under such alternative parameter regime, we performed similar analysis as the five different scenarios in Fig. 4A-B. The results are shown in Fig. 4C-D. In simulations of a network with intact long-range connections and increased local excitatory connections (in Fig. 4D and also in Fig. 4F), we changed the long-range connections strength *μ*_*EE*_ = 0.19*nA*. In Fig. 4C, when we gradually decrease the inhibitory connection strength *g*_*EI*,0_ from 0.5*nA* to 0 (from blue dots to orange dots), analytical calculations demonstrate that the stable low firing rate state disappears via a saddle node bifurcation at *g*_*EI*,0_ = 0.175*nA* (for area AIp). This demonstrates that, upon removal of inhibition, the high firing rate in Fig. 4C corresponds to a distinct state and not simply a shift of the baseline state.

In the increased local excitatory connection regime, we further introduced temporary external input to each local brain areas and record its stable firing rate shown in Fig. 4E.

In the simulation of Fig. 4F, we used the classic simulation protocol: an temporary external input is given to primary visual cortex and the delay period firing rate of each areas are recorded and shown.

### Thalamocortical network model

#### Corticothalamic connectivity

We introduced thalamic areas in the network to examine their effect on cortical dynamics. Each thalamic area includes 2 excitatory populations, A and B, with no inhibitory population. These two populations share the same selectivity with the corresponding cortical areas. Unlike cortical areas, there are no recurrent connections between thalamic neurons (Sherman 2007). Thalamic currents have the following contributions (tc stands for thalamocortical connections and ct for corticothalamic connections):

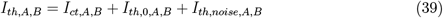

where *I*_*th,i*_ (*i* = *A, B*) is the total current received by each thalamic population, *I*_*ct,i*_ (*i* = *A, B*) is the long range current from cortical areas to target thalamic area, *I*_*th*,0,*i*_ (*i* = *A, B*) is the background current for each population, and *I*_*th,noise,i*_ (*i* = *A, B*) is the noise input to thalamic population A and B, which we set to 0 in our simulations. *I*_*ct,i*_ (*i* = *A, B*) has the following form:

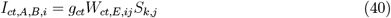

where *W*_*ct,E,ij*_ is the LR connectivity to thalamic neurons, and *S*_*k,j*_ is the synaptic variable of population *k* (*k* = *A, B*) in cortical area *j*. Since all thalamic neurons are excitatory, we model corticothalamic projections as in the previous section:

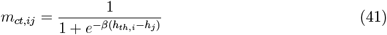

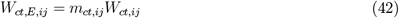

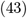

where

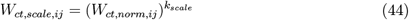

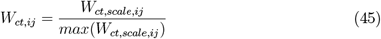

*W*_*ct,norm,ij*_ is the normalized connection strength from cortical area j to thalamic area i. *m*_*ct,ij*_ is the coefficient quantifying how the long range connections target excitatory neurons based on cortical hierarchy *h*_*j*_ and thalamic hierarchy *h*_*th,i*_.

The thalamic firing rates are described by:

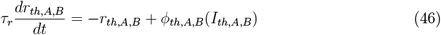

with the activation function for thalamic neurons given by:

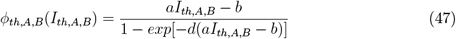

Thalamic neurons are described by AMPA synaptic variables (Jaramillo et al. 2019):

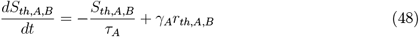

#### Thalamocortical connectivity

The connections from thalamic neurons to cortical neurons follow these equations

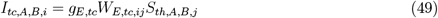

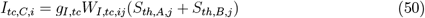

and connectivity

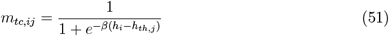

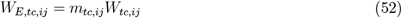

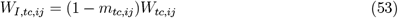

and connectivity matrix

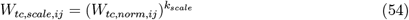

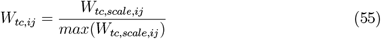

The thalamocortical input is added to the total input current of each cortical population.

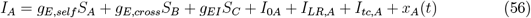

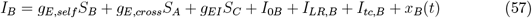

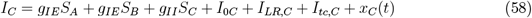

### Calculation of network structural measures

We considered three types of structural measures. The first one is input strength. Input strength of area *i* is the summation of the connection strengths onto node *i*. It quantifies the total external input onto area *i*.

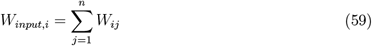

The second one is eigenvector centrality (Newman 2018). Eigenvector centrality of area *i* is the ith element of the leading eigenvector of the connectivity matrix. It quantifies how many areas are connected with the target area *i* and how important these neighbors are. *W* is a matrix where each element is *W*_*ij*_.

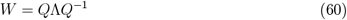

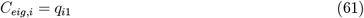

The third structural measure is loop strength, which quantifies how each area is involved in strong recurrent loops. We first define the strength of a single loop *k*

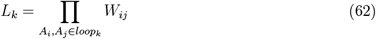

and then the loop strength 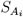 of a single area *A*_*i*_

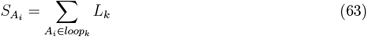

We now focus on cell type-specific structural measures. Cell type specificity is introduced via a coefficient *k*_*cell*_ that scales all long range connection strengths (cell type projection coefficient):

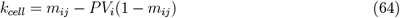

Thus, we can define cell type-specific connectivity as:

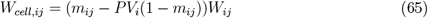

The cell type-specific connectivity is further normalized so that the maximum value is 1.

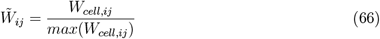

and cell type-specific input strength could be defined as:

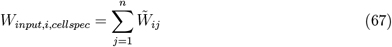

Similarly, cell type-specific eigenvector centrality is defined as

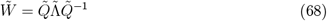

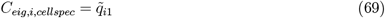

where 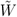 is a matrix where each element is 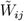 and the cell type-specific loop strength is defined as:

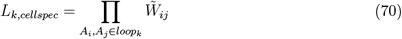

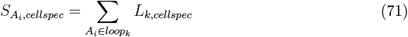

As a comparison, we also calculated the sign-only loop strength and no PV loop strength. We can define sign-only connectivity as:

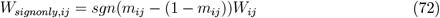

where sgn(x) is the sign function, which returns positive or negative values based on the sign of x. The major difference between sign only connectivity and cell type specific connectivity is that the strength of long range projection bias are not considered except the sign of it.We can also define no-PV connectivity as:

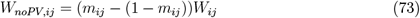

The difference between no-PV connectivity and cell-type specific connectivity is that the different strengths of local connections for each area are not considered in the no-PV connectivity.

We also used the sign-only and no-PV variants of connectivity measures to predict the delay period firing rate and classify core areas. This enabled us to compare these simplified measures to the cell-type-specific connectivity measures.

### Stimulation protocol and inhibition analysis

The model is simulated using an stochastic differential equation solver: Euler-Maruyama method. We write customized program using Python to implement this numerical method. The time step is set to be *dt*, and all the firing rates, synaptic variables and currents are initialized to be zero.

We simulate a working memory task by applying an external current *I*_*stim*_ to one of the excitatory populations, which represents a sensory (e.g., visual) stimulus that is to be remembered across a delay period. The external current is a pulsed input with start time *T*_*on*_ and offset time *T*_*off*_. Without losing generality, we assume that the external input is provided to population A. In most of the simulations in this study, we simulate a visual working memory task, with the external input applied to VISp. The simulation duration is *T*_*trial*_ and we used a time step of *dt*. The delay period is defined as the duration between the offset time *T*_*off*_ and trial end *T*_*trial*_. In order to obtain a stable firing rate, the delay period firing rate is calculated by averaging the firing rate from 2 seconds until the end of the delay period to 0.5 seconds until end. Firing rate, PV cell fraction, and hierarchy are plotted on a 3d brain surface using the website scalable brain atlas (https://scalablebrainatlas.incf.org/index.php). We apply inhibition analysis to understand the robustness of attractors and, more importantly, to investigate which areas play an important role in maintaining the attractor state. Excitatory input was applied to the inhibitory population I to simulate opto-genetic inhibition. The external input *I*_*inh*_ is strong as compared to *I*_*stim*_ and results in an elevated firing rate of the inhibitory population, which in turn decreases the firing rate of the excitatory populations. Usually the inhibition is applied to a single area. When inhibition is applied during the stimulus period, its start and end times are equal to *T*_*on*_ and *T*_*off*_, respectively. When inhibition is applied during delay period, its start time is later than *T*_*off*_ to allow the system settle to a stable state. Thus, the onset of inhibition starts 2 seconds after *T*_*off*_ and lasts until the end of trial. In the case of thalamocortical network simulations, we inhibit thalamic areas by introducing a hyperpolarizng current to both excitatory populations, since we do not have inhibitory populations in thalamic areas in the model.

To quantify the effect of single area or multiple areas inhibition, we calculate the average firing rate of areas that satisfy two conditions: i) the area shows persistent activity before inhibition and ii) the area does not receive inhibitory input. The ratio between such average firing rate after inhibition and before inhibition is used to quantify the overall effect of inhibition. If the ratio is lower than 100%, this suggests that inhibiting certain area(s) disrupts the maintenance of the attractor state. Note that the inhibition effect is typically not very strong, and only in rare cases, inhibition of a single area leads to loss of activity of other areas (Fig. 7B, Fig. 7C). To quantify such differences, we use a threshold of 10% to differentiate them. We will use (relatively) “weak inhibition effect” and “strong inhibition effect” to refer to them afterwards.

We used the three measures to classify areas into 4 types (Fig. 7D): i) inhibition effect during delay period, ii) inhibition effect during stimulus period, and iii) delay period firing rate. Areas with strong inhibition effect during stimulus period are classified as input areas; areas with strong inhibition effect during delay period and strong delay period firing rate are classified as core areas; areas with weak inhibition effect during delay period but strong firing rate are classified as readout areas; areas with weak inhibition effect during delay period and weak firing rate during delay period are classified as nonessential areas.

As an extension of the single area inhibition study, we focus on the role of readout areas. A pair of readout areas is randomly chosen and inhibited during the delay period under a similar protocol as the single area inhibition study. The inhibition effect, i.e., the decrement of the delay period firing rates of other non inhibited areas, is first quantified for each inhibition pair (*A*_*i*_, *A*_*j*_). Next, the inhibition effect is averaged one more time for each area *A*_*i*_ across all inhibition pairs that includes the area ((*A*_*i*_, *A*_*j*_), where *j* ≠ *i*). An anologous procedure is performed for triplets and quadruplets of readout areas. Additionally, we also calculate the mean inhibition effect between pair of areas, which are both selected from core areas, both selected from readout areas, or we chose one area from core areas, one area from readout areas.

### Simulation of multiple attractors

Multiple attractors coexist in the network and its properties and number depends on the connectivity and dynamics of each node. In this study we did not try to capture all the possible attractors in the network, but rather compare the number of attractors for different networks. Here we briefly describe the protocol used to identify multiple attractors in the network. We first choose k areas and then generate a subset of areas as the stimulation areas. We cover all possible subsets, which means we run 2^*k*^ simulations in total. The external stimulus is given to all areas in the subset simultaneously with same strength and duration. The delay period activity is then quantified using a similar protocol as the standard simulation protocol. The selection of k areas corresponds to a qualitative criterion. First we choose the areas with small PV fraction or high hierarchy, since these areas are more likely to show persistent activity. Second, the number of possible combination grows exponentially as we increase k, and if we use k = 43, the number of combinations is around 8.8e+12, which is beyond our simulation power. As a trade-off between the simulation power and coverage of areas, we choose k = 18, which correspond to 2.6e+5 different combinations of stimulation. For each parameter setting, we run 2.6e+5 simulations to capture possible attractor patterns. For each attractor pattern, a binary vector is generated by thresholding delay firing rate using a firing rate threshold of 5Hz. An attractor pattern is considered distinct if and only if the binary vector is different from all identified attractors. In these way we can identify different attractors in the simulation. We also apply same simulation pipeline to identify attractors for different parameters. Specifically we change the long range connectivity strength *μ*_*EE*_ and local excitatory connections *g*_*E,self*_.

## Data availability

The manuscript constitutes a computational study, so no experimental data has been generated. The simulation and analysis code will be available in GitHub upon publication.

---

**Figure 1 - Supplement 1.**
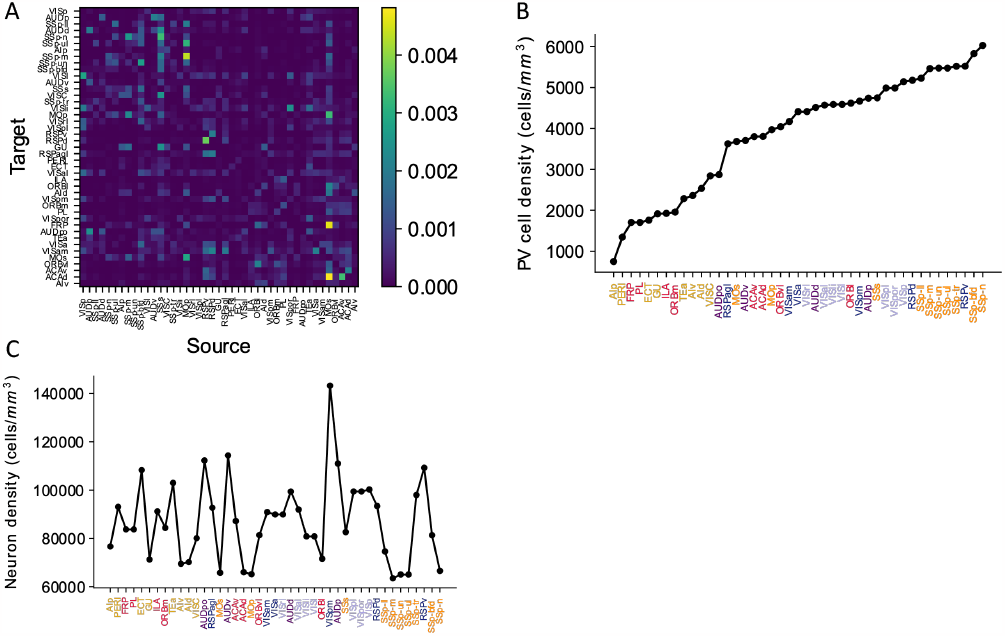
Anatomical details of the mouse cortex. (A). Connectivity matrix depicting cortico-cortical connections between 43 cortical areas. Areas are sorted according to their hierarchy. (B). The raw PV cell density for each cortical area (Y axis), with areas sorted (X axis). Each area belongs to one of five modules, shown in color (see also Fig. 1). (Harris et al. 2019). (C). Neuron density for each cortical area with same sorted order as (B). The data is from Erö et al. 2018.

**Figure 2 - Supplement 1.**
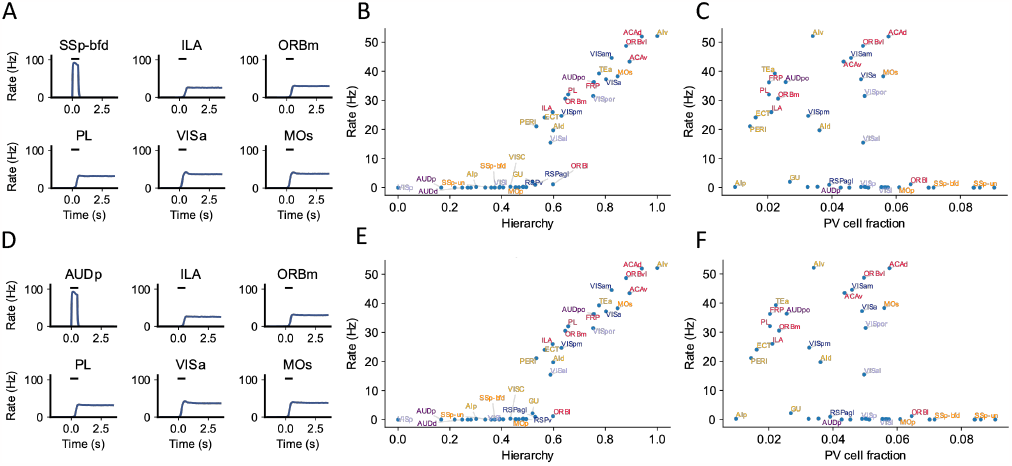
Example simulation for different sensory modalities. The simulation protocol is the same as the default one in Fig. 2, except that the external input is applied to primary sensory areas related to two other sensory modalities: somatosensory and auditory. (A). The activity of 6 selected areas during the working memory task is shown. A somatosensory input of 500ms is applied to primary somatosensory area SSp-bfd, which propagates to the rest of the large-scale network. (B). Similar to the simulation where a primary visual area is stimulated (Fig. 2D), delay period firing for somatosensory stimulation is positively correlated with cortical hierarchy (r = 0.89, p<0.05). (C). Delay period firing rate is moderately correlated with PV cell fraction (r =−0.4, p<0.05). (D),(E) and (F) are similar to (A), (B) and (C) except that the input is given to primary auditory area AUDp. (E). Delay period firing is also positively correlated with cortical hierarchy (r = 0.89, p<0.05). (F). Delay period firing rate is moderately correlated with PV cell fraction (r = −0.4, p<0.05)

**Figure 3 - Supplement 1.**
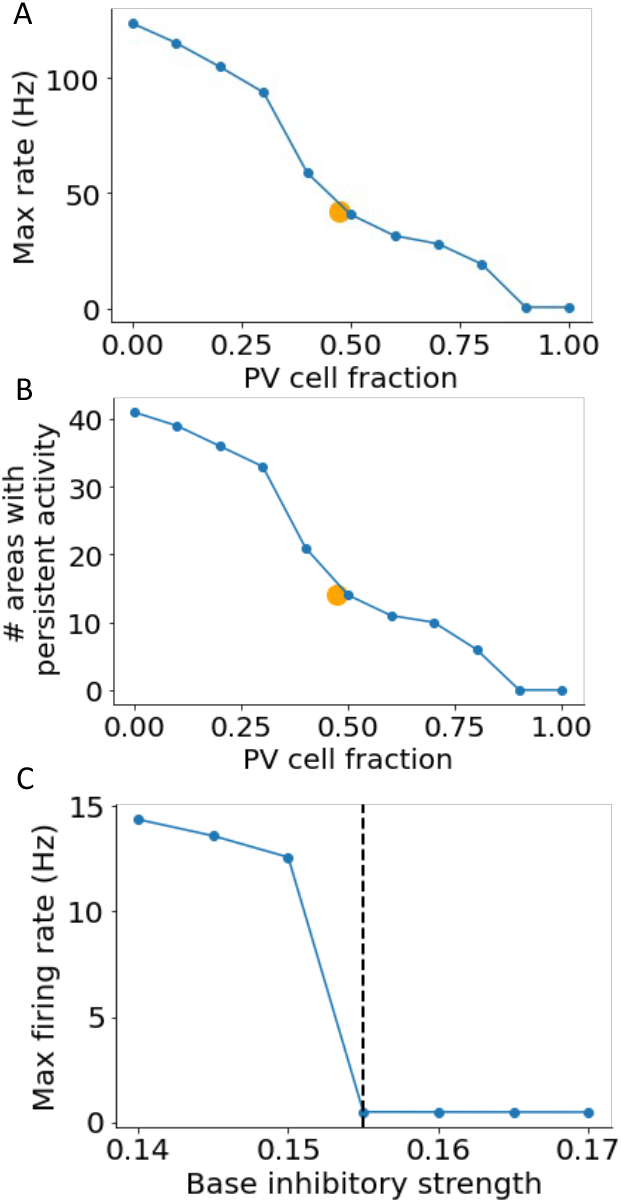
Dependence of persistent activity on inhibitory model parameters (A). The maximum firing rate of all areas depends on the constant PV cell fraction in models without a gradient of PV. Average PV cell fraction from the anatomical data is shown as an orange dot. (B). Same as (A), except for the number of areas showing persistent activity. (C). Firing rate during the delay period for local circuits without long-range projections as a function of base inhibitory strength. If the base inhibitory strength is larger than a threshold (0.155, marked by the dashed line), none of the areas show independent persistent activity.

**Figure 5 - Supplement 1.**
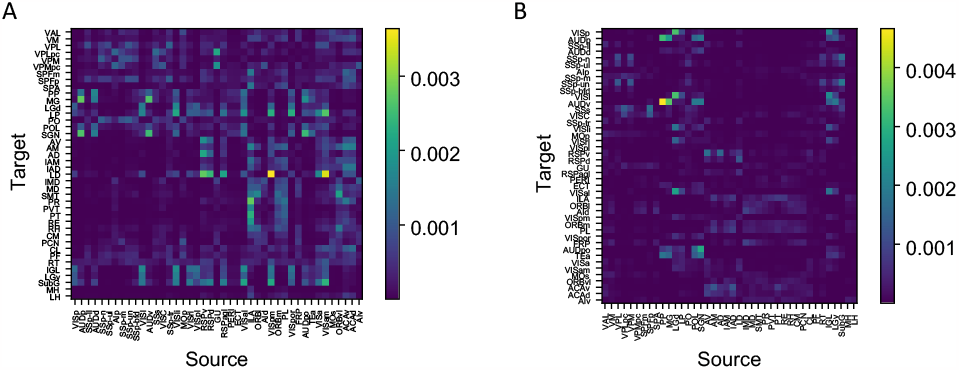
Anatomical data of thalamus and cortical connectivity. (A). Connectivity matrix of corticothalamic connections: 43 cortical areas to 40 thalamic areas. (B). Connectivity matrix of thalamocortical connections: 40 thalamic areas to 43 cortical areas.

**Figure 6 - Supplement 1.**
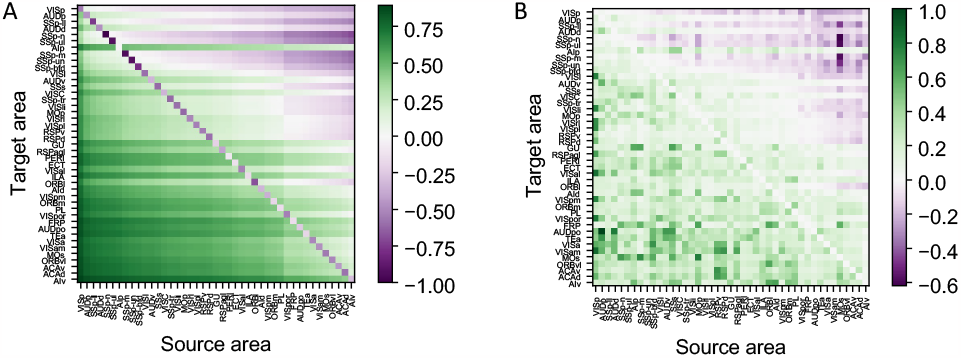
Details of cell type-specific connectivity measures. (A). The matrix of cell type projection coefficients between cortical areas. The cell type projection coefficient is given by the formula k_cell_ = m_ij_−PV_i_(1−m_ij_). (B). The matrix of connectivity strengths, modified by cell type projection coefficient between cortical areas. The modified connectivity strength is given by 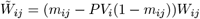.

**Figure 6 - Supplement 2.**
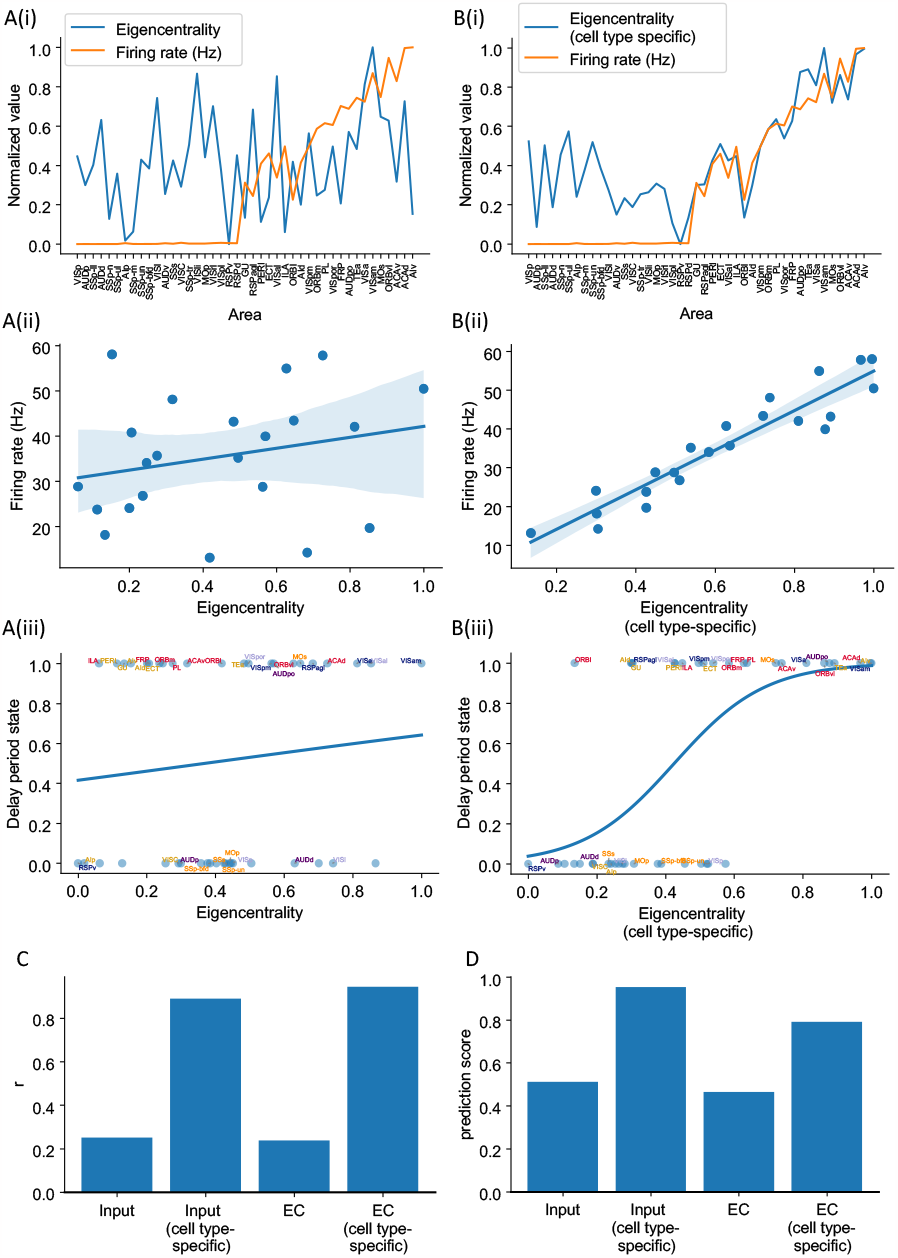
Cell type-specific eigenvector centrality measures are better at predicting firing rate patterns than raw eigenvector centrality measures. The analysis is the same as in Fig. 6, where we compared cell type-specific input strength and raw input strength. Eigenvector centrality (EC, eigencentrality) of area i is the ith element of the leading eigenvector of the connectivity matrix. It quantifies how many areas are connected with the target area i and how important are these neighbors. Details are in the Methods section. (A(i)). Delay period firing rate (orange) and eigenvector centrality for each cortical area (blue). (A(ii)). Eigenvector centrality does not show a significant correlation with delay period firing rate for areas showing persistent activity in the model (r = 0.24, p = 0.29). (A(iii)). Eigenvector centrality cannot be used to predict whether an area shows persistent activity or not (prediction accuracy = 0.46). (B(i)). Delay period firing rate (orange) and cell type-specific eigenvector centrality for each cortical area (blue). (B(ii)). Cell type-specific eigenvector centrality has a strong correlation with the firing rate of cortical areas showing persistent activity (r = 0.94, p < 0.05). (B(iii)). Cell type-specific eigenvector centrality predicts whether an area shows persistent activity or not (prediction accuracy = 0.79). (C). Comparison of the correlation coefficient r for raw eigenvector centrality and cell type-specific eigenvector centrality in predicting delay firing rate. Raw input strength and cell type-specific input strength are also included for comparison. (D). Comparison of the prediction accuracy for raw eigenvector centrality and cell type-specific eigenvector centrality. Raw input strength and cell type-specific input strength are also included for comparison.

**Figure 6 - Supplement 3.**
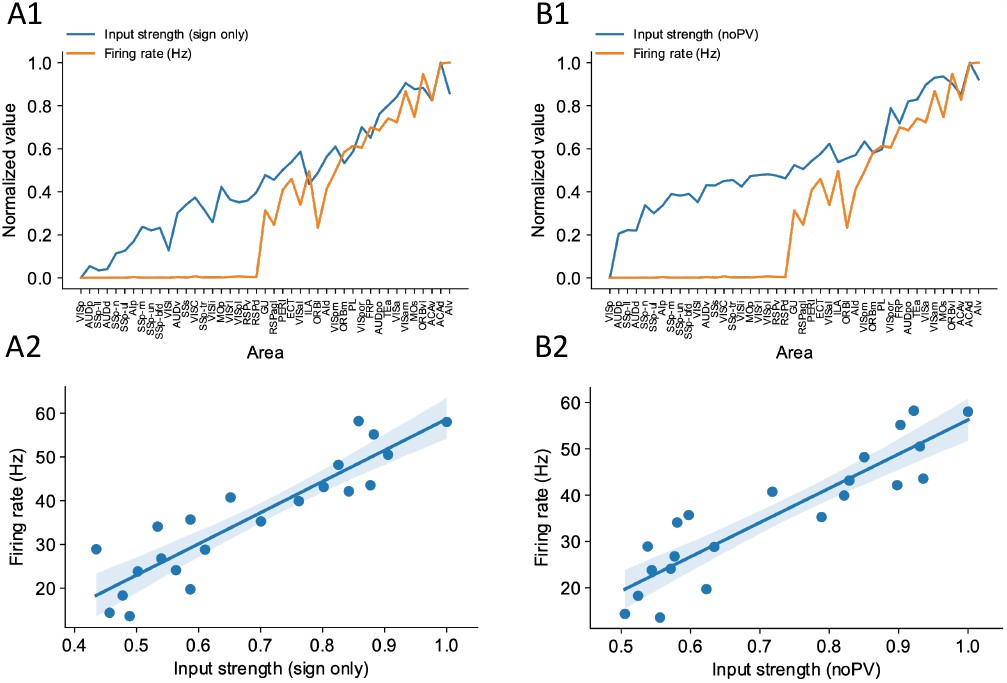
Sign only input strength measure and noPV input strength measure predict firing rate well. (A1). Delay period firing rate (orange) and sign only input strength for each cortical areas. (A2). Sign only input strength has a strong correlation with delay period firing rate of cortical areas showing persistent activity. (r =0.90, p <0.05) (B1). Delay period firing rate (orange) and noPV input strength for each cortical areas. (B2). noPV input strength has a strong correlation with delay period firing rate of cortical areas showing persistent activity (r =0.90, p <0.05).

**Figure 7 - Supplement 1.**
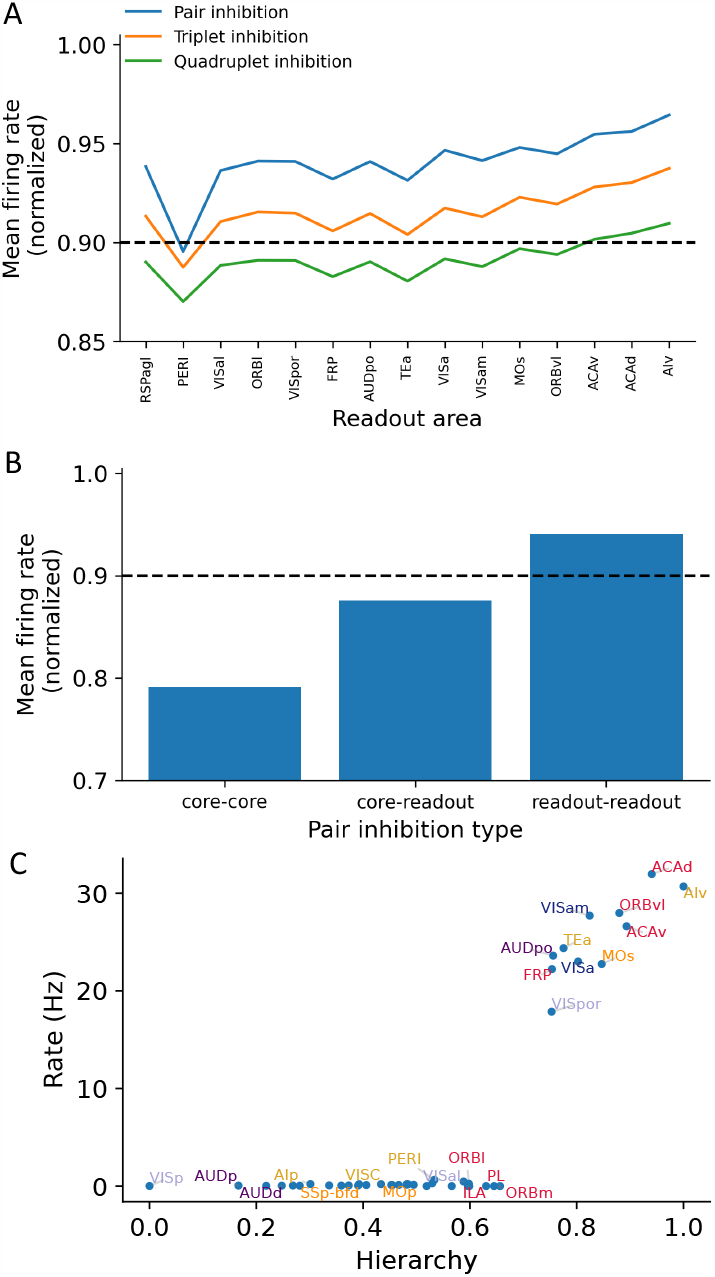
Multiple-area inhibition experiments demonstrate the relative importance for core and readout areas in maintaining network-level persistent activity. (A). The x-axis shows readout areas that are inhibited as part of a pair (blue), triplet (orange), or quadruplet (green). For any given readout area A, the y-axis shows the average firing rate of all cortical areas that exhibit persistent activity when A was inhibited as part of the inhibited pair (triplet, quadruplet). The decrement in delay period activity is stronger as more areas are inhibited. (B). Bar plots showing the average firing rate of the network after inhibition of pair-wise combinations of core and readout areas. For example, the bar plot for ‘readout-readout’ is the average firing rate for all readout-readout areas pairs and is corresponding to the blue curve in (A). Dashed line in (A) and (B) denotes a threshold below which we consider an ‘inhibition effect’ to be significant. (C). Delay period firing rates as a function of hierarchy after inhibition of all core areas during the delay period. Although some readout areas show persistent firing, there is 48% decrement in average firing rate.

**Figure 8 - Supplement 1.**
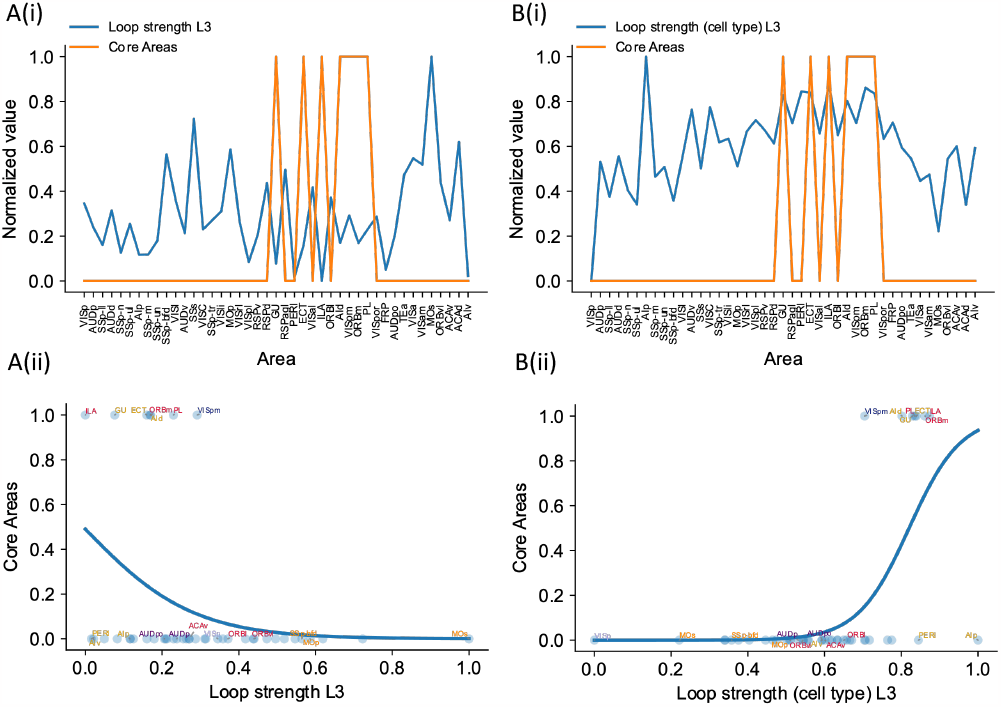
Cell type-specific loop strengths (Length 3 loops) are also better at predicting firing rate patterns than raw loop measures. Loop strengths (length 3 loops or L3) is calculated using similar method as loop strengths (length 2 loops). The only difference is we considered loops with length 3 (eg. A1->A2->A3->A1). The analysis is the same as in Fig. 7, where we compared cell type-specific loop strengths (length 2 loops) and raw loop strengths. (A(i)). Loop strength (blue) is plotted alongside Core Areas (orange), a binary variable that takes the value 1 if the area is indeed a Core Area, 0 otherwise. Loop strength is normalized to a range of (0, 1) for better comparison. (A(ii)). A high loop strength value does not imply that an area is a Core Area. (B(i)). Same as (A), but for cell type-specific loop strength. (B(ii)). High cell type-specific loop measures predicts that an area is a Core Area (prediction accuracy = 0.95). Same as (A), but for cell type-specific loop strength.

**Figure 8 - Supplement 2.**
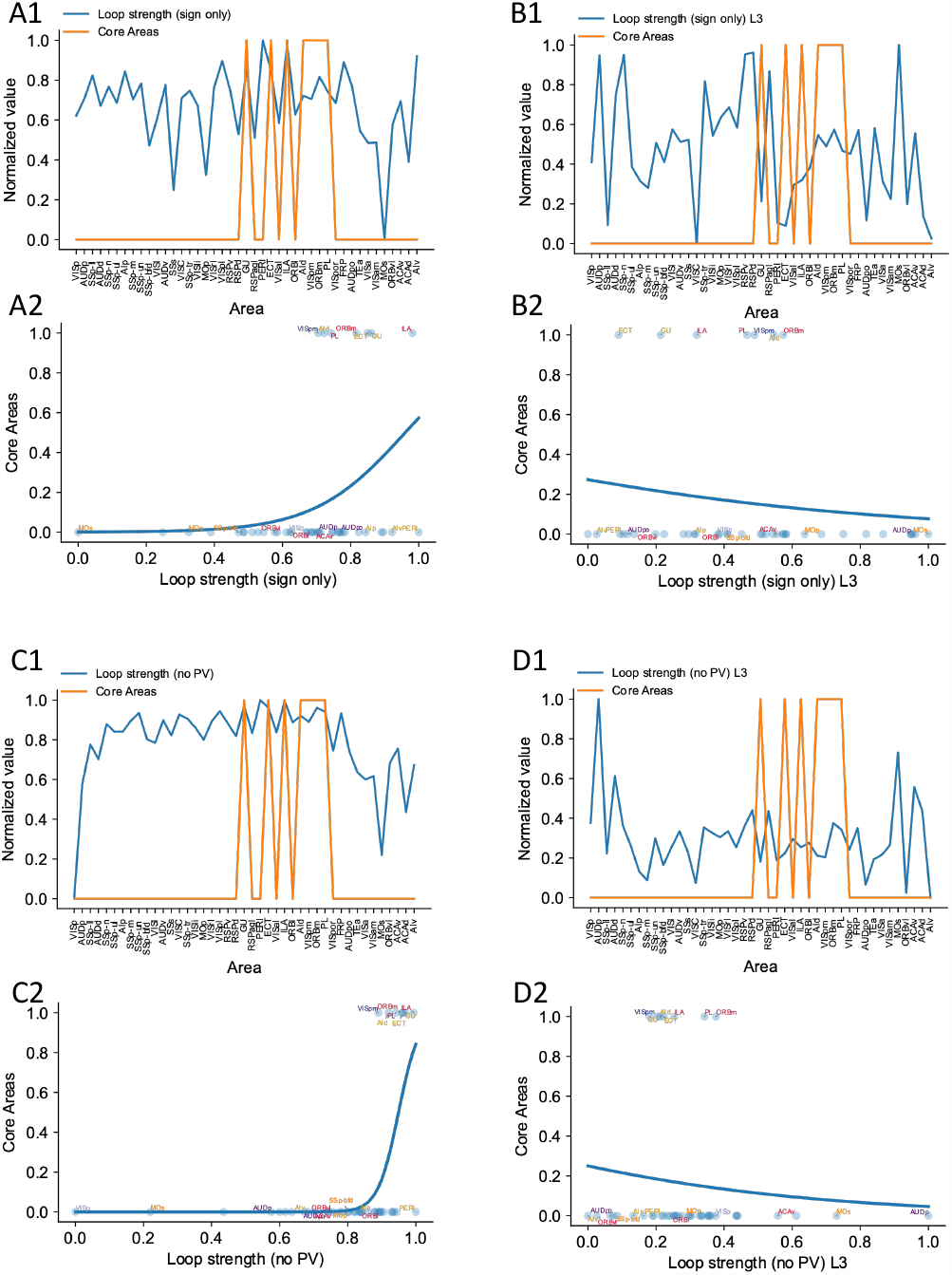
(A1). Relationship between core areas (orange) and length 2 (sign only) loop strength. Areas are sorted according to their hierarchy. Whether an area is a core area is represented in either 0 or 1. (A2). High loop strength is a good predictor of whether an area is a core area. Blue curve shows the logistic regression analysis used to differentiate the core areas versus non core areas (prediction accuracy = 0.83). (B1) and (B2). Same as (A1) and (A2), but with length 3 sign only loop strength. Length 3 sign only loop strength does not show a positive relationship with core areas (prediction accuracy = 0.83) (C1). (C2). Same as (A1) and (A2), except for comparing whether an area is a core area (orange) and length 2 noPV loop strength. Length 2 noPV loop strength predicts the core areas. prediction accuracy = 0.90 (D1). (D2). Same to A1 and A2, except for comparing whether an area is a core area (orange) and length 3 noPV loop strength. Length 3 noPV loop strength does not show a positive relationship with core areas. prediction accuracy = 0.83.

**Table 1:** Supplementary experimental evidence. The listed literature include experiments that provide supporting evidence for working memory activity in cortical and subcortical brain areas in the mouse or rat. These studies show either that a given area is involved in working memory tasks and/or exhibit delay period activity. Area name corresponds to what has been reported in the literature. Some areas do not correspond exactly to the names from the Allen common coordinate framework.

| Area | Supporting literature |
| --- | --- |
| ALM (MOs) | (Kopec et al. 2015; Guo et al. 2014; Li et al. 2016b),<br>(Inagaki et al. 2019; Erlich et al. 2011; Guo et al. 2017),<br>(Gilad et al. 2018; Gao et al. 2018),<br>(Wu et al. 2020; Voitov and Mrsic-Flogel 2022) |
| mPFC (PL/ILA) | (Liu et al. 2014; Schmitt et al. 2017),<br>(Bolkan et al. 2017) |
| OFC | (Wu et al. 2020) |
| PPC (VISa, VISam) | (Harvey et al. 2012; Voitov and Mrsic-Flogel 2022) |
| AIa (AId, AIv) | (Zhu et al. 2020) |
| Area p (VISpl) | (Gilad et al. 2018) |
| dorsal cortex | (Pinto et al. 2019) |
| entorhinal (in vitro persistent activity) | (Egorov et al. 2002) |
| piriform | (Zhang et al. 2019; Wu et al. 2020) |
| VM/VAL | (Guo et al. 2017) |
| MD | (Schmitt et al. 2017; Bolkan et al. 2017) |
| superior colliculus | (Kopec et al. 2015) |
| cerebellar nucleus | (Gao et al. 2018) |

**Table 2:**
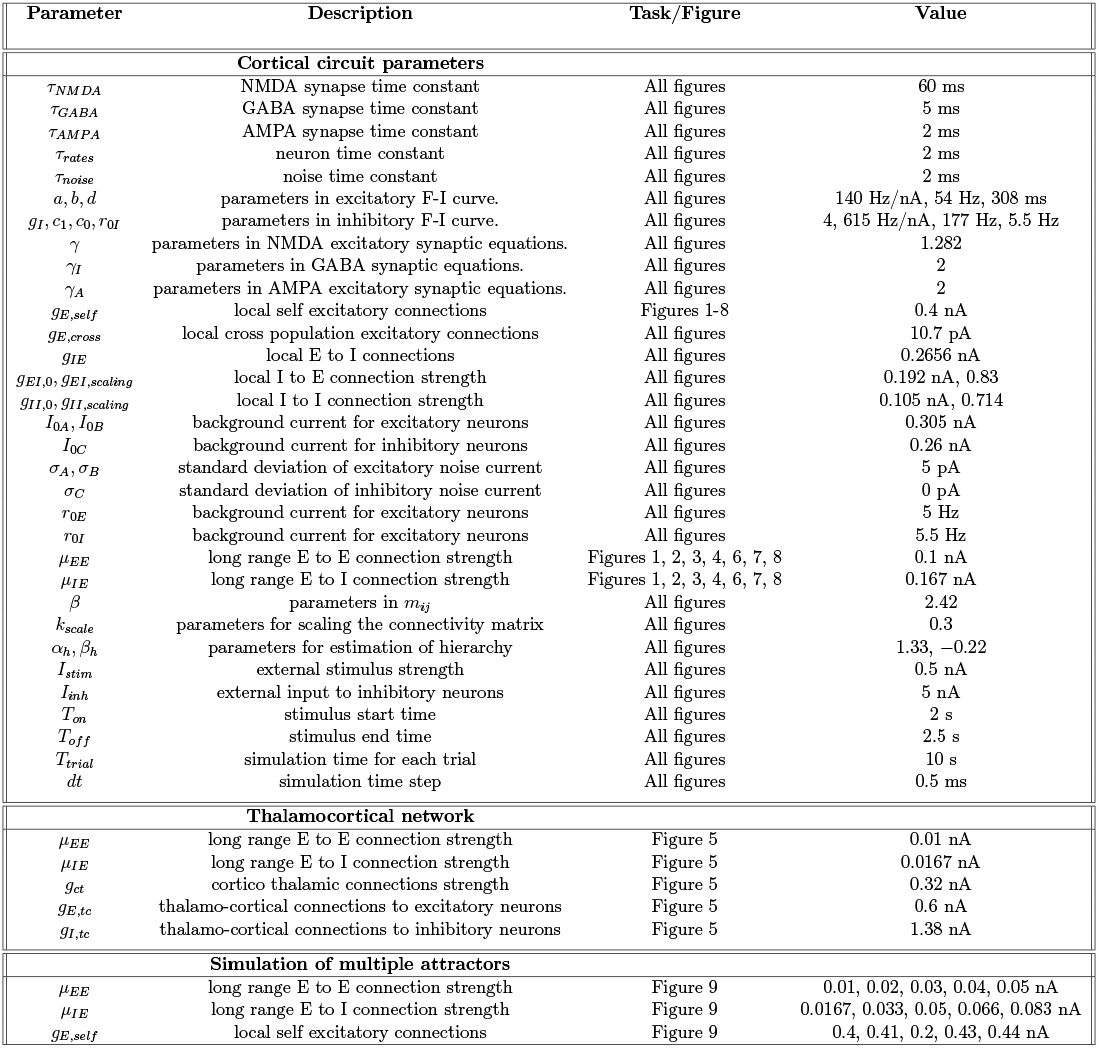
Parameters for numerical simulations.

## References

Abbott, L. F. et al. (2020). “The mind of a mouse”. Cell 182, pp. 1372–1376.

Lorente de Nó, R. (1933). “Vestibulo-ocular reflex arc”. Arch. Neurol. Psych. 30, pp. 245–291.

Abbott, Larry F. et al. (2017). “An International Laboratory for Systems and Computational Neuroscience”. Neuron 96.6, pp. 1213–1218.

Baddeley, Alan (2012). “Working Memory: Theories, Models, and Controversies.” Annual review of psychology 63.1, pp. 1–29. pmid: 21961947.

Ballesteros-Yáñez, Inmaculada, Ruth Benavides-Piccione, Jean-Pierre Bourgeois, Jean-Pierre Changeux, and Javier DeFelipe (2010). “Alterations of Cortical Pyramidal Neurons in Mice Lacking High-Affinity Nicotinic Receptors”. Proceedings of the National Academy of Sciences 107.25, pp. 11567–11572.

Bassett, Danielle S. and Edward T. Bullmore (2017). “Small-World Brain Networks Revisited”. The Neuroscientist 23.5, pp. 499–516.

Bolkan, Scott S., Joseph M. Stujenske, Sebastien Parnaudeau, Timothy J. Spellman, Caroline Rauffenbart, Atheir I. Abbas, Alexander Z. Harris, Joshua A. Gordon, and Christoph Kellendonk (2017). “Thalamic Projections Sustain Prefrontal Activity during Working Memory Maintenance”. Nature Neuroscience 20.7 (7), pp. 987–996.

Burt, Joshua B., Murat Demirtaş, William J. Eckner, Natasha M. Navejar, Jie Lisa Ji, William J. Martin, Alberto Bernacchia, Alan Anticevic, and John D. Murray (2018). “Hierarchy of Transcriptomic Specialization across Human Cortex Captured by Structural Neuroimaging Topography”. Nature Neuroscience 21.9, pp. 1251–1259.

Cabral, Joana, Etienne Hugues, Olaf Sporns, and Gustavo Deco (2011). “Role of Local Network Oscillations in Resting-State Functional Connectivity”. NeuroImage 57.1, pp. 130–139.

Campagnola, Luke et al. (2022). “Local Connectivity and Synaptic Dynamics in Mouse and Human Neocortex”. Science 375.6585, eabj5861.

Chaudhuri, Rishidev, Kenneth Knoblauch, Marie-Alice Gariel, Henry Kennedy, and Xiao-Jing Wang (2015). “A Large-Scale Circuit Mechanism for Hierarchical Dynamical Processing in the Primate Cortex”. Neuron 88.2, pp. 419–431.

Christophel, Thomas B., P. Christiaan Klink, Bernhard Spitzer, Pieter R. Roelfsema, and John-Dylan Haynes (2017). “The Distributed Nature of Working Memory”. Trends in Cognitive Sciences 21.2, pp. 111–124.

Crabtree, John W. (2018). “Functional Diversity of Thalamic Reticular Subnetworks”. Frontiers in Systems Neuroscience 12, p. 41.

D. J. Amit (1995). “The Hebbian paradigm reintegrated: local reverberations as internal representations”. Behav. Brain Sci. 18, pp. 617–626.

Deco, Gustavo, Adrián Ponce-Alvarez, Patric Hagmann, Gian Luca Romani, Dante Mantini, and Maurizio Corbetta (2014). “How Local Excitation–Inhibition Ratio Impacts the Whole Brain Dynamics”. Journal of Neuroscience 34.23, pp. 7886–7898. pmid: 24899711.

Demirtaş, Murat, Joshua B. Burt, Markus Helmer, Jie Lisa Ji, Brendan D. Adkinson, Matthew F. Glasser, David C. Van Essen, Stamatios N. Sotiropoulos, Alan Anticevic, and John D. Murray (2019). “Hierarchical Heterogeneity across Human Cortex Shapes Large-Scale Neural Dynamics”. Neuron 101.6, 1181–1194.e13.

Diesmann, Markus, Marc-Oliver Gewaltig, and Ad Aertsen (1999). “Stable Propagation of Synchronous Spiking in Cortical Neural Networks”. Nature 402.6761 (6761), pp. 529–533.

Dotson, Nicholas M., Steven J. Hoffman, Baldwin Goodell, and Charles M. Gray (2018). “Feature-Based Visual Short-Term Memory Is Widely Distributed and Hierarchically Organized”. Neuron 99.1, 215–226.e4.

Egorov, Alexei V., Bassam N. Hamam, Erik Fransén, Michael E. Hasselmo, and Angel A. Alonso (2002). “Graded Persistent Activity in Entorhinal Cortex Neurons”. Nature 420.6912, pp. 173–178.

Elston, Guy N. (2007). “Specialization of the Neocortical Pyramidal Cell during Primate Evolution”. In: Evolution of Nervous Systems. Elsevier, pp. 191–242.

Elston, Guy N. and Marcello G.P. Rosa (1998). “Complex Dendritic Fields of Pyramidal Cells in the Frontal Eye Field of the Macaque Monkey: Comparison with Parietal Areas 7a and LIP”. NeuroReport 9.1, pp. 127–131.

Erlich, Jeffrey C., Max Bialek, and Carlos D. Brody (2011). “A Cortical Substrate for Memory-Guided Orienting in the Rat”. Neuron 72.2, pp. 330–343.

Erö, Csaba, Marc-Oliver Gewaltig, Daniel Keller, and Henry Markram (2018). “A Cell Atlas for the Mouse Brain”. Frontiers in Neuroinformatics 12, p. 84.

Felleman, D J and D C Van Essen (1991). “Distributed Hierarchical Processing in the Primate Cerebral Cortex.” Cerebral Cortex 1.1, pp. 1–47. pmid: 1822724.

Froudist-Walsh, Sean, Daniel P. Bliss, Xingyu Ding, Lucija Rapan, Meiqi Niu, Kenneth Knoblauch, Karl Zilles, Henry Kennedy, Nicola Palomero-Gallagher, and Xiao-Jing Wang (2021). “A Dopamine Gradient Controls Access to Distributed Working Memory in the Large-Scale Monkey Cortex”. Neuron 109.21, 3500–3520.e13.

Froudist-Walsh, Sean, Ting Xu, Meiqi Niu, Lucija Rapan, Ling Zhao, Daniel S. Margulies, Karl Zilles, Xiao-Jing Wang, and Nicola Palomero-Gallagher (2023). “Gradients of Neurotransmitter Receptor Expression in the Macaque Cortex”. Nature Neuroscience 26.7, pp. 1281–1294.

Fulcher, Ben D., John D. Murray, Valerio Zerbi, and Xiao-Jing Wang (2019). “Multimodal Gradients across Mouse Cortex”. Proceedings of the National Academy of Sciences 116.10, pp. 4689–4695. pmid: 30782826.

Funahashi, Shintaro, Charles J. Bruce, and Patricia S. Goldman-Rakic (1989). “Mnemonic Coding of Visual Space in the Monkey’s Dorsolateral Prefrontal Cortex”. Journal of Neurophysiology 61.2, pp. 331–349.

Fuster, Joaquin M. and Garrett E. Alexander (1971). “Neuron Activity Related to Short-Term Memory”. Science 173.3997, pp. 652–654. pmid: 4998337.

Gămănuţ, Răzvan, Henry Kennedy, Zoltán Toroczkai, Mária Ercsey-Ravasz, David C. Van Essen, Kenneth Knoblauch, and Andreas Burkhalter (2018). “The Mouse Cortical Connectome, Characterized by an Ultra-Dense Cortical Graph, Maintains Specificity by Distinct Connectivity Profiles”. Neuron 97.3, 698–715.e10.

Gao, Zhenyu, Courtney Davis, Alyse M Thomas, Michael N Economo, Amada M Abrego, Karel Svoboda, Chris I De Zeeuw, and Nuo Li (2018). “A Cortico-Cerebellar Loop for Motor Planning.” Nature 39, p. 1062. pmid: 30333626.

Gilad, Ariel, Yasir Gallero-Salas, Dominik Groos, and Fritjof Helmchen (2018). “Behavioral Strategy Determines Frontal or Posterior Location of Short-Term Memory in Neocortex”. Neuron 99.4, 814–828.e7.

Gilman, Joshua P., Maria Medalla, and Jennifer I. Luebke (2017). “Area-Specific Features of Pyramidal Neurons—a Comparative Study in Mouse and Rhesus Monkey”. Cerebral Cortex 27.3, pp. 2078–2094.

Goldman-Rakic, P. S (1995). “Cellular Basis of Working Memory”. Neuron 14.3, pp. 477–485.

Guo, Zengcai V., Hidehiko K. Inagaki, Kayvon Daie, Shaul Druckmann, Charles R. Gerfen, and Karel Svoboda (2017). “Maintenance of Persistent Activity in a Frontal Thalamocortical Loop”. Nature 545.7653 (7653), pp. 181–186.

Guo, Zengcai V., Nuo Li, Daniel Huber, Eran Ophir, Diego Gutnisky, Jonathan T. Ting, Guoping Feng, and Karel Svoboda (2014). “Flow of Cortical Activity Underlying a Tactile Decision in Mice”. Neuron 81.1, pp. 179–194.

Hádinger, Nóra, Emília Bősz, Boglárka Tóth, Gil Vantomme, Anita Lüthi, and László Acsády (2023). “Region-Selective Control of the Thalamic Reticular Nucleus via Cortical Layer 5 Pyramidal Cells”. Nature Neuroscience 26.1, pp. 116–130.

Harris, Julie A. et al. (2019). “Hierarchical Organization of Cortical and Thalamic Connectivity”. Nature 575.7781 (7781), pp. 195–202.

Harvey, Christopher D., Philip Coen, and David W. Tank (2012). “Choice-Specific Sequences in Parietal Cortex during a Virtual-Navigation Decision Task”. Nature 484.7392, pp. 62–68. pmid: 22419153.

Honey, Christopher J., Rolf Kötter, Michael Breakspear, and Olaf Sporns (2007). “Network Structure of Cerebral Cortex Shapes Functional Connectivity on Multiple Time Scales”. Proceedings of the National Academy of Sciences 104.24, pp. 10240–10245. pmid: 17548818.

Inagaki, Hidehiko K., Lorenzo Fontolan, Sandro Romani, and Karel Svoboda (2019). “Discrete Attractor Dynamics Underlies Persistent Activity in the Frontal Cortex”. Nature 566.7743 (7743), pp. 212–217.

Inagaki, Hidehiko K., Miho Inagaki, Sandro Romani, and Karel Svoboda (2018). “Low-Dimensional and Monotonic Preparatory Activity in Mouse Anterior Lateral Motor Cortex”. The Journal of Neuroscience 38.17, pp. 4163–4185.

Jaramillo, Jorge, Jorge F. Mejias, and Xiao-Jing Wang (2019). “Engagement of Pulvino-cortical Feedforward and Feedback Pathways in Cognitive Computations”. Neuron 101.2, 321–336.e9.

Javadzadeh, Mitra and Sonja B. Hofer (2022). “Dynamic Causal Communication Channels between Neocortical Areas”. Neuron 0.0.

Joglekar, Madhura R., Jorge F. Mejias, Guangyu Robert Yang, and Xiao-Jing Wang (2018). “Inter-Areal Balanced Amplification Enhances Signal Propagation in a Large-Scale Circuit Model of the Primate Cortex”. Neuron 98.1, 222–234.e8. pmid: 29576389.

Jones, Edward G. (2007). “Neuroanatomy: Cajal and after Cajal”. Brain Research Reviews 55.2, pp. 248–255.

Jonikaitis, Donatas, Behrad Noudoost, and Tirin Moore (2023). Dissociating the Contributions of Frontal Eye Field Activity to Spatial Working Memory and Motor Preparation. preprint. Neuroscience.

Jun, James J. et al. (2017). “Fully Integrated Silicon Probes for High-Density Recording of Neural Activity”. Nature 551.7679 (7679), pp. 232–236.

Kim, Yongsoo, Guangyu Robert Yang, Kith Pradhan, Kannan Umadevi Venkataraju, Mihail Bota, Luis Carlos García del Molino, Greg Fitzgerald, Keerthi Ram, Miao He, Jesse Maurica Levine, Partha Mitra, Z. Josh Huang, Xiao-Jing Wang, and Pavel Osten (2017). “Brain-Wide Maps Reveal Stereotyped Cell-Type-Based Cortical Architecture and Subcortical Sexual Dimorphism”. Cell 171.2, 456–469.e22.

Klatzmann, Ulysse, Sean Froudist-Walsh, Daniel P. Bliss, Panagiota Theodoni, Jorge F. Mejias, Meiqi Niu, Lucija Rapan, Nicola Palomero-Gallagher, Claire Sergent, Stanislas Dehaene, and Xiao-Jing Wang (2022). “A Connectome-Based Model of Conscious Access in Monkey Cortex”. bioRxiv, p. 2022.02.20.481230.

Knox, Joseph E., Kameron Decker Harris, Nile Graddis, Jennifer D. Whitesell, Hongkui Zeng, Julie A. Harris, Eric Shea-Brown, and Stefan Mihalas (2018). “High-Resolution Data-Driven Model of the Mouse Connectome”. Network Neuroscience 3.1, pp. 217–236. pmid: 30793081.

Kopec, Charles D., Jeffrey C. Erlich, Bingni W. Brunton, Karl Deisseroth, and Carlos D. Brody (2015). “Cortical and Subcortical Contributions to Short-Term Memory for Orienting Movements”. Neuron 88.2, pp. 367–377. pmid: 26439529.

Kubanek, Jan and Lawrence H. Snyder (2015). “Reward-Based Decision Signals in Parietal Cortex Are Partially Embodied”. The Journal of Neuroscience 35.12, pp. 4869–4881.

Leavitt, Matthew L., Diego Mendoza-Halliday, and Julio C. Martinez-Trujillo (2017). “Sustained Activity Encoding Working Memories: Not Fully Distributed”. Trends in Neurosciences 40.6, pp. 328–346.

Li, Nuo, Tsai-Wen Chen, Zengcai V. Guo, Charles R. Gerfen, and Karel Svoboda (2015). “A Motor Cortex Circuit for Motor Planning and Movement”. Nature 519.7541 (7541), pp. 51–56.

Li, Nuo, Kayvon Daie, Karel Svoboda, and Shaul Druckmann (2016a). “Robust Neuronal Dynamics in Premotor Cortex during Motor Planning”. Nature 532.7600, pp. 459–464.

Li, Nuo, Kayvon Daie, Karel Svoboda, and Shaul Druckmann (2016b). “Robust Neuronal Dynamics in Premotor Cortex during Motor Planning”. Nature 532.7600, pp. 459–464.

Liu, Ding, Xiaowei Gu, Jia Zhu, Xiaoxing Zhang, Zhe Han, Wenjun Yan, Qi Cheng, Jiang Hao, Hongmei Fan, Ruiqing Hou, Zhaoqin Chen, Yulei Chen, and Chengyu T. Li (2014). “Medial Prefrontal Activity during Delay Period Contributes to Learning of a Working Memory Task”. Science 346.6208, pp. 458–463.

Markov, Nikola T, Julien Vezoli, Pascal Chameau, Arnaud Falchier, René Quilodran, Cyril Huissoud, Camille Lamy, Pierre Misery, Pascale Giroud, Shimon Ullman, Pascal Barone, Colette Dehay, Kenneth Knoblauch, and Henry Kennedy (2014a). “Anatomy of Hierarchy: Feedforward and Feedback Pathways in Macaque Visual Cortex.” Journal of Comparative Neurology 522.1, pp. 225–259. pmid: 23983048.

Markov, Nikola T. et al. (2014b). “A Weighted and Directed Interareal Connectivity Matrix for Macaque Cerebral Cortex”. Cerebral Cortex 24.1, pp. 17–36.

Mejias, Jorge F., John D. Murray, Henry Kennedy, and Xiao-Jing Wang (2016). “Feedforward and Feedback Frequency-Dependent Interactions in a Large-Scale Laminar Network of the Primate Cortex”. Science Advances 2.11, e1601335.

Mejias, Jorge F. and Xiao-Jing Wang (2022). “Mechanisms of Distributed Working Memory in a Large-Scale Network of Macaque Neocortex”. eLife 11, e72136.

Meng, John Hongyu, Benjamin Schuman, Bernardo Rudy, and Xiao-Jing Wang (2023). “Mechanisms of Dominant Electrophysiological Features of Four Subtypes of Layer 1 Interneurons”. The Journal of Neuroscience 43.18, pp. 3202–3218.

Murphy, B. K. and K. D. Miller (2009). “Balanced amplification: a new mechanism of selective amplification of neural activity patterns”. Neuron 61, pp. 635–648.

Murray, John D, Alberto Bernacchia, David J Freedman, Ranulfo Romo, Jonathan D Wallis, Xinying Cai, Camillo Padoa-Schioppa, Tatiana Pasternak, Hyojung Seo, Daeyeol Lee, and Xiao-Jing Wang (2014). “A Hierarchy of Intrinsic Timescales across Primate Cortex”. Nature Neuroscience 17.12, pp. 1661–1663.

Murray, John D., Jorge Jaramillo, and Xiao-Jing Wang (2017). “Working Memory and Decision-Making in a Frontoparietal Circuit Model”. Journal of Neuroscience 37.50, pp. 12167–12186. pmid: 29114071.

Musall, S., M. T. Kaufman, A. L. Juavinett, S. Gluf, and A. K. Churchland (2019). “Single-trial neural dynamics are dominated by richly varied movements”. Nat. Neurosci. 22, pp. 1677–1686.

Myme, Chaelon I. O., Ken Sugino, Gina G. Turrigiano, and Sacha B. Nelson (2003). “The NMDA-to-AMPA Ratio at Synapses Onto Layer 2/3 Pyramidal Neurons Is Conserved Across Prefrontal and Visual Cortices”. Journal of Neurophysiology 90.2, pp. 771–779.

Naskar, Shovan, Jia Qi, Francisco Pereira, Charles R. Gerfen, and Soohyun Lee (2021). “Cell-Type-Specific Recruitment of GABAergic Interneurons in the Primary Somatosensory Cortex by Long-Range Inputs”. Cell Reports 34.8, p. 108774.

Newman, M. E. J. (2018). Networks. Second edition. Oxford: Oxford University Press. Nigro, Maximiliano José, Kasper Kjelsberg, Laura Convertino, Rajeevkumar Raveendran

Nair, and Menno P. Witter (2022). Enrichment of Specific GABAergic Neuronal Types in the Mouse Perirhinal Cortex. preprint. Neuroscience.

Oh, Seung Wook et al. (2014). “A Mesoscale Connectome of the Mouse Brain”. Nature 508.7495 (7495), pp. 207–214.

P. S. Goldman-Rakic (1995). “Cellular basis of working memory”. Neuron 14, pp. 477–485.

Petreanu, Leopoldo, Tianyi Mao, Scott M Sternson, and Karel Svoboda (2009). “The Subcellular Organization of Neocortical Excitatory Connections.” Nature 457.7233, pp. 1142–1145. pmid: 19151697.

Pinto, Lucas, Kanaka Rajan, Brian DePasquale, Stephan Y. Thiberge, David W. Tank, and Carlos D. Brody (2019). “Task-Dependent Changes in the Large-Scale Dynamics and Necessity of Cortical Regions”. Neuron 104.4, 810–824.e9.

R. J. Douglas, C. Koch, M. Mahowald, K. M. Martin, and H. H. Suarez (1995). “Recurrent excitation in neocortical circuits”. Science 269, pp. 981–985.

Sanzeni, Alessandro, Bradley Akitake, Hannah C Goldbach, Caitlin E Leedy, Nicolas Brunel, and Mark H Histed (2020). “Inhibition Stabilization Is a Widespread Property of Cortical Networks”. eLife 9, e54875.

Schmidt, Maximilian, Rembrandt Bakker, Kelly Shen, Gleb Bezgin, Markus Diesmann, and Sacha Jennifer van Albada (2018). “A Multi-Scale Layer-Resolved Spiking Network Model of Resting-State Dynamics in Macaque Visual Cortical Areas”. PLOS Computational Biology 14.10, e1006359.

Schmitt, L. Ian, Ralf D. Wimmer, Miho Nakajima, Michael Happ, Sima Mofakham, and Michael M. Halassa (2017). “Thalamic Amplification of Cortical Connectivity Sustains Attentional Control”. Nature 545.7653 (7653), pp. 219–223.

Shen, Shan, Xiaolong Jiang, Federico Scala, Jiakun Fu, Paul Fahey, Dmitry Kobak, Zhenghuan Tan, Na Zhou, Jacob Reimer, Fabian Sinz, and Andreas S. Tolias (2022). “Distinct Organization of Two Cortico-Cortical Feedback Pathways”. Nature Communications 13.1, p. 6389.

Sherman, S Murray (2007). “The Thalamus Is More than Just a Relay”. Current opinion in neurobiology 17.4, pp. 417–422.

Shine, James M, Matthew J Aburn, Michael Breakspear, and Russell A Poldrack (2018). “The Modulation of Neural Gain Facilitates a Transition between Functional Segregation and Integration in the Brain”. eLife 7. Ed. by Gustavo Deco, e31130.

Siu, Caitlin, Justin Balsor, Sam Merlin, Frederick Federer, and Alessandra Angelucci (2021). “A Direct Interareal Feedback-to-Feedforward Circuit in Primate Visual Cortex”. Nature Communications 12.1, p. 4911.

Steinmetz, Nicholas A., Peter Zatka-Haas, Matteo Carandini, and Kenneth D. Harris (2019). “Distributed Coding of Choice, Action and Engagement across the Mouse Brain”. Nature 576.7786 (7786), pp. 266–273.

Steinmetz, Nicholas A. et al. (2021). “Neuropixels 2.0: A Miniaturized High-Density Probe for Stable, Long-Term Brain Recordings”. Science 372.6539, eabf4588.

Stringer, Carsen, Marius Pachitariu, Nicholas Steinmetz, Charu Bai Reddy, Matteo Carandini, and Kenneth D. Harris (2019). “Spontaneous Behaviors Drive Multidimensional, Brainwide Activity”. Science 364.6437, eaav7893.

Suzuki, Mototaka and Jacqueline Gottlieb (2013). “Distinct Neural Mechanisms of Distractor Suppression in the Frontal and Parietal Lobe”. Nature Neuroscience 16.1 (1), pp. 98–104.

Svoboda, Karel and Nuo Li (2018). “Neural Mechanisms of Movement Planning: Motor Cortex and Beyond”. Current Opinion in Neurobiology 49, pp. 33–41.

Torres-Gomez, Santiago, Jackson D Blonde, Diego Mendoza-Halliday, Eric Kuebler, Michelle Everest, Xiao Jing Wang, Wataru Inoue, Michael O Poulter, and Julio Martinez-Trujillo (2020). “Changes in the Proportion of Inhibitory Interneuron Types from Sensory to Executive Areas of the Primate Neocortex: Implications for the Origins of Working Memory Representations”. Cerebral Cortex 30.8, pp. 4544–4562.

Tremblay, Robin, Soohyun Lee, and Bernardo Rudy (2016). “GABAergic Interneurons in the Neocortex: From Cellular Properties to Circuits”. Neuron 91.2, pp. 260–292.

Tsodyks, Misha V., William E. Skaggs, Terrence J. Sejnowski, and Bruce L. McNaughton (1997). “Paradoxical Effects of External Modulation of Inhibitory Interneurons”. The Journal of Neuroscience 17.11, pp. 4382–4388.

Vincis, Roberto, Ke Chen, Lindsey Czarnecki, John Chen, and Alfredo Fontanini (2020). “Dynamic Representation of Taste-Related Decisions in the Gustatory Insular Cortex of Mice”. Current Biology 30.10, 1834–1844.e5.

Voitov, Ivan and Thomas D. Mrsic-Flogel (2022). “Cortical Feedback Loops Bind Distributed Representations of Working Memory”. Nature, pp. 1–9.

Wang, Peng, Ru Kong, Xiaolu Kong, Raphaël Liégeois, Csaba Orban, Gustavo Deco, Martijn P. van den Heuvel, and B. T. Thomas Yeo (2019). “Inversion of a Large-Scale Circuit Model Reveals a Cortical Hierarchy in the Dynamic Resting Human Brain”. Science Advances 5.1, eaat7854.

Wang, Quanxin et al. (2020). “The Allen Mouse Brain Common Coordinate Framework: A 3D Reference Atlas”. Cell 181.4, 936–953.e20. pmid: 32386544.

Wang, X.-J. (2002). “Probabilistic decision making by slow reverberation in cortical circuits”. Neuron 36, pp. 955–968.

Wang, X.-J., J. Tegnér, C. Constantinidis, and P. S. Goldman-Rakic (2004). “Division of Labor among Distinct Subtypes of Inhibitory Neurons in a Cortical Microcircuit of Working Memory”. Proceedings of the National Academy of Sciences 101.5, pp. 1368–1373.

Wang, Xiao-Jing (1999). “Synaptic Basis of Cortical Persistent Activity: The Importance of NMDA Receptors to Working Memory”. Journal of Neuroscience 19.21, pp. 9587–9603.

Wang, Xiao-Jing (2001). “Synaptic Reverberation Underlying Mnemonic Persistent Activity”. Trends in Neurosciences 24.8, pp. 455–463.

Wang, Xiao-Jing (2020). “Macroscopic Gradients of Synaptic Excitation and Inhibition in the Neocortex”. Nature Reviews Neuroscience 21.3, pp. 169–178.

Wang, Xiao-Jing (2021). “50 years of mnemonic persistent activity: Quo vadis?” Trends in Neurosci. 44, pp. 888–902.

Wang, Xiao-Jing (2022). “Theory of the Multiregional Neocortex: Large-Scale Neural Dynamics and Distributed Cognition”. Annual Review of Neuroscience 45.1, pp. 533–560.

Wang, Yu, Xinxin Yin, Zhouzhou Zhang, Jiejue Li, Wenyu Zhao, and Zengcai V. Guo (2021). “A Cortico-Basal Ganglia-Thalamo-Cortical Channel Underlying Short-Term Memory”. Neuron 109.21, 3486–3499.e7.

Wildenberg, Gregg A., Matt R. Rosen, Jack Lundell, Dawn Paukner, David J. Freedman, and Narayanan Kasthuri (2021). “Primate Neuronal Connections Are Sparse in Cortex as Compared to Mouse”. Cell Reports 36.11, p. 109709.

Wong, Kong-Fatt and Xiao-Jing Wang (2006). “A Recurrent Network Mechanism of Time Integration in Perceptual Decisions”. Journal of Neuroscience 26.4, pp. 1314–1328. pmid: 16436619.

Wu, Zheng, Ashok Litwin-Kumar, Philip Shamash, Alexei Taylor, Richard Axel, and Michael N. Shadlen (2020). “Context-Dependent Decision Making in a Premotor Circuit”. Neuron 106.2, 316–328.e6.

Xu, Yaoda (2017). “Reevaluating the Sensory Account of Visual Working Memory Storage”. Trends in Cognitive Sciences 21.10, pp. 794–815.

Yang, Guangyu Robert, Madhura R. Joglekar, H. Francis Song, William T. Newsome, and Xiao-Jing Wang (2019). “Task Representations in Neural Networks Trained to Perform Many Cognitive Tasks”. Nature Neuroscience 22.2, pp. 297–306.

Yang, Sheng-Tao, Min Wang, Constantinos D Paspalas, Johanna L Crimins, Marcus T Altman, James A Mazer, and Amy F T Arnsten (2018). “Core Differences in Synaptic Signaling Between Primary Visual and Dorsolateral Prefrontal Cortex”. Cerebral Cortex 28.4, pp. 1458–1471.

Yizhar, Ofer, Lief E. Fenno, Thomas J. Davidson, Murtaza Mogri, and Karl Deisseroth (2011). “Optogenetics in Neural Systems”. Neuron 71.1, pp. 9–34.

Zandvakili, Amin and Adam Kohn (2015). “Coordinated Neuronal Activity Enhances Cortic-ocortical Communication.” Neuron 87.4, pp. 827–839. pmid: 26291164.

Zhang, Xiaoxing, Wenjun Yan, Wenliang Wang, Hongmei Fan, Ruiqing Hou, Yulei Chen, Zhaoqin Chen, Chaofan Ge, Shumin Duan, Albert Compte, and Chengyu T Li (2019). “Active Information Maintenance in Working Memory by a Sensory Cortex”. eLife 8, e43191.

Zhu, Jia, Qi Cheng, Yulei Chen, Hongmei Fan, Zhe Han, Ruiqing Hou, Zhaoqin Chen, and Chengyu T. Li (2020). “Transient Delay-Period Activity of Agranular Insular Cortex Controls Working Memory Maintenance in Learning Novel Tasks”. Neuron 105.5, 934– 946.e5.

